# Flexible integration of corollary discharge and sensory feedback signals in somatosensory cortex

**DOI:** 10.64898/2026.04.02.716126

**Authors:** Xinyue An, Raeed H. Chowdhury, Kyle P. Blum, Lee E. Miller, Joshua I. Glaser

**Affiliations:** Department of Neurology, Northwestern University, Chicago, IL, USA; Interdepartmental Neuroscience Program, Northwestern University, Chicago, IL, USA; Department of Bioengineering, University of Pittsburgh, Pittsburgh, PA, USA; Center for the Neural Basis of Cognition, University of Pittsburgh, Pittsburgh, PA, USA; Department of Neuroscience, Northwestern University, Chicago, IL, USA; Department of Physical Medicine and Rehabilitation, Northwestern University, Chicago, IL, USA; Shirley Ryan AbilityLab, Chicago, IL, USA; Department of Biomedical Engineering, Northwestern University, Evanston, IL, USA; Department of Computer Science, Northwestern University, Evanston, IL, USA; National Institute for Theory and Mathematics in Biology, Chicago, IL, USA

## Abstract

Motor control depends on the continuous integration of motor and sensory signals to maintain accurate estimates of body state, yet neural evidence for this integration remains elusive. Here, we investigated the interaction of motor corollary discharge and proprioceptive feedback signals in area 2 of monkey somatosensory cortex during voluntary and externally-perturbed reaching tasks. Though single neurons had mixed responses to corollary discharge and sensory feedback, we disentangled these signals at the population level to discover they occupy approximately orthogonal subspaces. Integrating information across these subspaces enabled accurate body state estimation prior to feedback arrival during voluntary movements. Moreover, the orthogonal population geometry of corollary discharge and sensory feedback enabled cancellation of movement-related signals to improve the decoding of external perturbations. Together, these results identified orthogonality as a population-level coding strategy for flexible integration of motor and sensory signals to support multiple distinct computations.

## Main

Accurate estimation of body state is essential for guiding ongoing movements^1–5^. A key mechanism supporting this process is corollary discharge—internal copies of motor commands routed to sensory systems and cerebellum^6–9^. Such signaling is central to optimal feedback control theory, which proposes that the sensory system integrates corollary discharge with incoming sensory feedback to construct internal estimates of body state^10–13^ (**Figure 1A**). Corollary discharge allows predictions of expected sensory input, which, in the Bayesian framework of optimal feedback control, act as priors that can then be combined with incoming, delayed feedback (“likelihood”) to form an optimal estimate of body state (“posterior”)^14–17^. Despite ample theoretical work, neural evidence for this integration remains limited.

**Figure 1.**
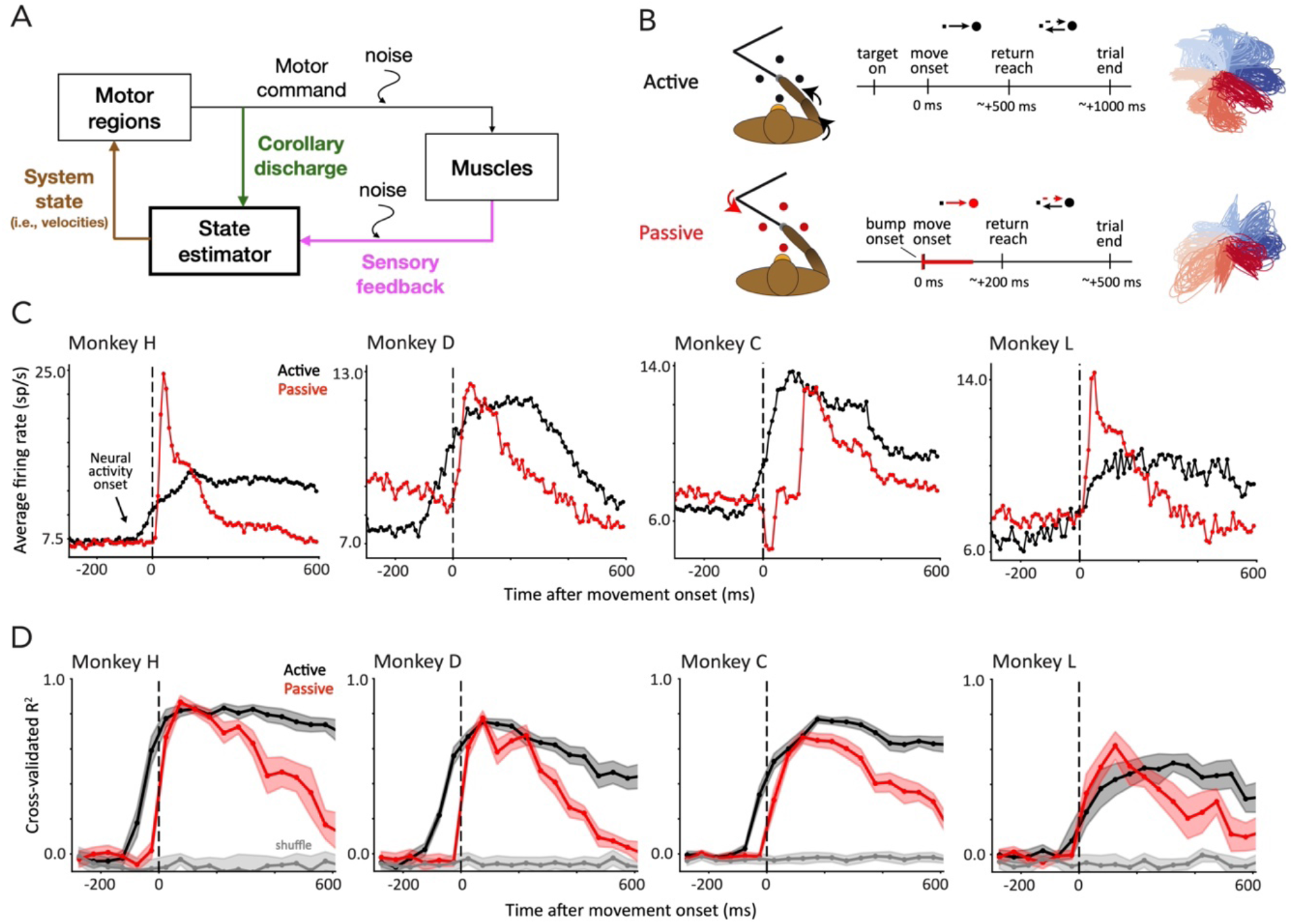
Area 2 receives corollary discharge signals. **A**. Schematic of Optimal Feedback Control theory. The sensory system acts as a state estimator that integrates corollary discharge and sensory feedback signals to form body state estimates. **B**. Active and passive reaching tasks. *Left*: Trial structure. Black segments denote voluntary movements; red segments denote externally perturbed movements. *Right*: Hand trajectories across trials in Monkey H, colored by reach direction. **C**. Average neural activity onset precedes movement onset in active trials. **D**. Pre-movement activity decodes upcoming reach directions. We used linear regression to decode reach directions in active trials (black), bump directions in passive trials (red), or their shuffled directions for control (gray). Decoding was performed in 50-ms windows centered at the plotted time. Shading shows standard deviation across 20 validation folds. Lines connect discrete time-bin estimates for visualization, which gives the appearance of a rise before time 0 in passive trials. Also note passive trials are aligned to movement, not bump, onset.

Somatosensory cortex, and in particular Brodmann’s area 2, is well positioned to support the integration of these signals. As a higher-order processing region within the somatosensory hierarchy^18^, area 2 receives convergent proprioceptive and cutaneous inputs and encodes high-level movement kinematics^19–23^. Complementing its sensory input, area 2 also receives input from motor cortex^18,24,25^—a likely route for corollary discharge signals. This is supported by physiological evidence that its neural responses can precede voluntary muscle activity and differ between active and passive movements^19,22,26–30^. With this combination of sensory and motor signals, area 2 may convey integrated information to downstream motor and parietal regions to support motor control and learning^24,25^. These features make area 2 a compelling site to investigate the neural basis of sensorimotor integration.

To understand how corollary discharge and somatosensory feedback signals interact, we recorded from area 2 in macaque monkeys during active and passive “center-out reaching” tasks. We first disentangled corollary discharge and sensory feedback signals within population activity based on their different onset times relative to active movements, and validated this decomposition using mechanical perturbations during passive trials. Intriguingly, the two signals occupied approximately orthogonal neural subspaces, enabling their simultaneous yet distinct population-level representations. Consistent with optimal feedback control theory, although each signal alone supported kinematic decoding at distinct time lags, their integration enabled accurate kinematic estimation prior to feedback arrival. Moreover, by analyzing the signals when reaching movements were mechanically perturbed during a “reach-bump” task, we discovered that orthogonality not only promotes state estimation during voluntary movements but also facilitates the detection of externally imposed movements. Thus, orthogonal population codes of corollary discharge and sensory feedback in area 2 provide a flexible scheme for dual functions of accurate state estimation and reliable perturbation detection.

## Results

We recorded neural spiking activity from four rhesus macaques (Monkeys H, D, C, and L) using multi-electrode arrays implanted in the arm representation of Brodmann’s area 2 in somatosensory cortex^19^ (**Extended Data Figure 1**). Each monkey was trained to use a manipulandum to complete an active-passive center-out task, where on each trial, the monkey either reached to a target presented on the screen, or was passively perturbed a short distance toward a target. In both trial types, the monkey actively returned to the center afterwards (**Figure 1B**). Monkeys H and D had eight targets (orthogonal and diagonal), while Monkeys C and L only had four orthogonal targets. The two trial types (active and passive) were randomly interleaved. Monkey D also completed reach-bump trials, where the monkey reached to a target as in an active trial, with a brief assistive or resistive perturbation (toward or against the target direction, respectively) delivered 250 ms after the go cue. We recorded the animal’s endpoint kinematics via manipulandum during the tasks.

### Corollary discharge signals predict future movements

We first investigated whether a corollary discharge signal was present in these recordings. We sought neural signals that (i) precede the sensory consequences of voluntary movement, distinguishing them from sensory feedback signals, and (ii) convey information about the upcoming movement, distinguishing them from the spontaneous or neuromodulatory signals that broadly activate the brain but lack movement-related details^31–33^. We aligned neural activity by movement onset times, and observed that, across animals, average neural activity increased before movement onset in active trials (**Figure 1C; Extended Data Figure 2A** for second sessions for Monkeys H, D). Significant differences from baseline activity emerged at 60, 100, 40, and 90 ms prior to movement for Monkeys H, D, C, L (p<0.05 via sliding-window t-test; **Methods**). This was not the case in passive trials (p>0.05 for times preceding movement onset). These latencies are consistent with postulated corollary discharge from motor cortex (M1), as it is slightly shorter than the typical ∼100 ms delay from M1 to movement onset (combining a ∼50 ms delay from M1 activity to EMG^34,35^ and ∼50 ms from EMG to movement^27,28^). Importantly, we could use this early neural activity to decode upcoming reach directions at above-chance levels (**Figure 1D**; **Extended Data Figure 2B**), consistent with a role in corollary discharge. We observed larger pre-movement increases in neural activity and substantially higher directional information in Monkeys H and D, suggesting stronger corollary discharge signals recorded in these animals. Accordingly, as our goal was investigating the integration of corollary discharge with feedback, we focused our subsequent analyses on data from Monkeys H and D (results from Monkeys C and L shown in **Extended Data** throughout).

**Figure 2.**
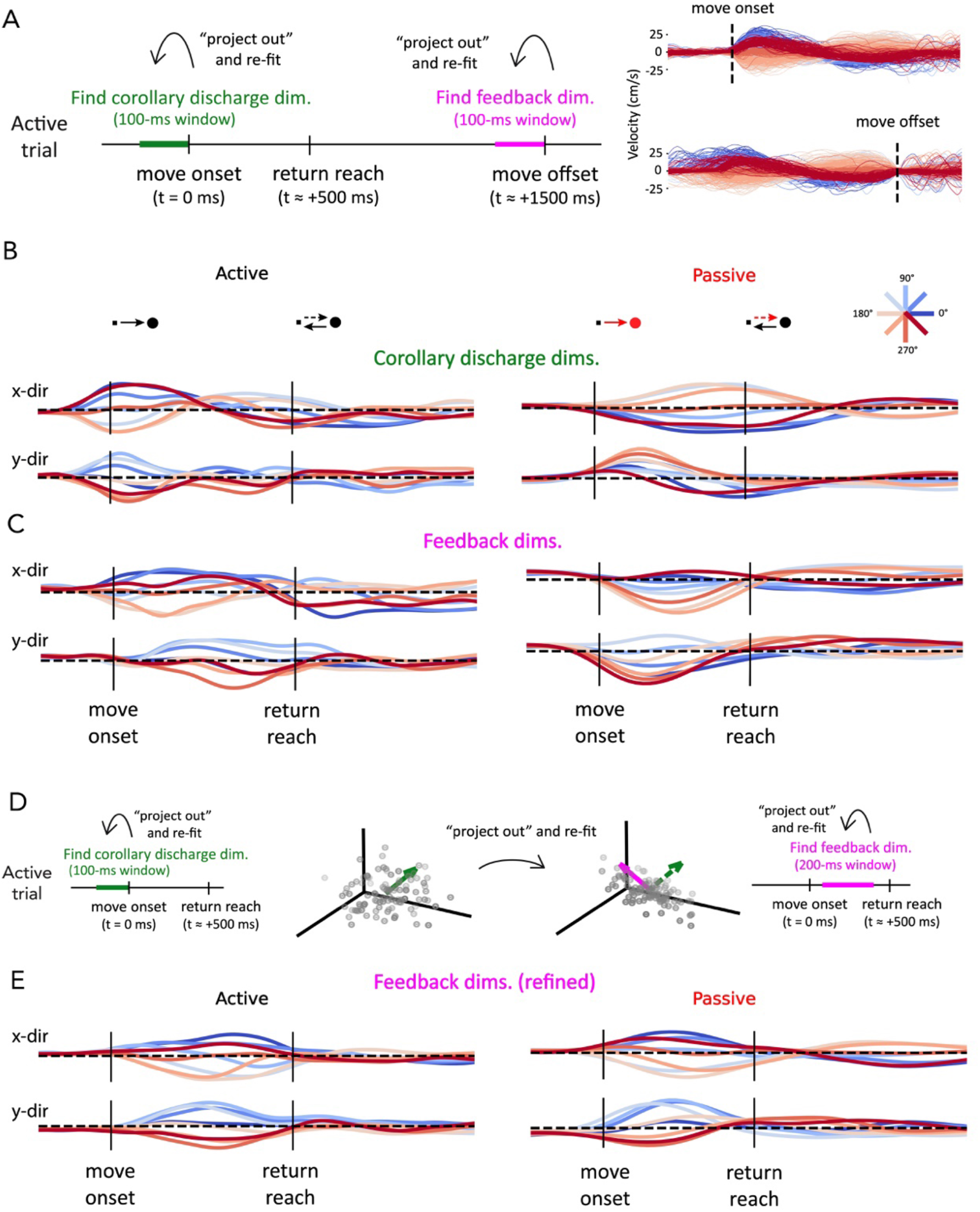
Extracting corollary discharge and feedback signals. **A**. Procedure for independent signal extraction. We extracted corollary-discharge and feedback signal subspaces within their respective time windows (*Left*). This requires aligning trials separately to movement onset or offset (*Right*). **B**. Neural activity of Monkey H projected onto the top corollary-discharge dimensions during Active (*Left*) and Passive (*Right*) trials, averaged across trials within each direction. Traces are colored by reach direction in Active trials and by bump direction in Passive trials. Plotted dimensions correspond to the first dimensions extracted for the x- and y-components of reach directions (i.e., the best decoder of those directions). **C**. Same as **B**, for the top feedback dimensions. These dimensions were extracted using neural activity during the return movement, which is opposite in direction to the reach used to extract corollary-discharge dimensions. Thus, for clearer visual comparison with **B**, we sign-flipped feedback dimensions during plotting. **D**. Procedure for stepwise signal extraction. We first extracted corollary discharge as in **A**, and then extracted feedback after removing corollary-discharge-related activity. **E**. Neural activity projected onto the top feedback dimensions obtained using the stepwise extraction procedure. Note that condition traces appear to diverge prior to movement onset due to smoothing, see **Extended Data Figure 5** for unsmoothed trajectories.

To characterize how corollary discharge is represented within the population activity, we aimed to identify a low-dimensional neural subspace to isolate the signal from other concurrent neural activity. We defined corollary discharge as neural activity prior to voluntary movement, which informs upcoming movement directions (is “direction-selective”). In the 100-ms window prior to movement onset, we found the signal’s constituent neural subspace by iteratively solving the linear regressions from neural activity to direction (sine and cosine) of the upcoming reach direction (**Figure 2A**, green). That is, we first fit a linear decoder, subtracted the projection of neural activity onto this decoder (which comprised one neural axis of the signal subspace) from the original activity, and then fit a new decoder to the residual activity (see **Methods**). We repeated the process iteratively until no directional information remained (held-out R^2^ ≤ 0). These axes comprised a multi-dimensional neural subspace of the signal (3-dimensional for both Monkeys H and D).

To understand the recovered corollary discharge dimensions, we first visualized their activity during active trials (**Figure 2B** left). As expected, direction-selective activity along corollary-discharge dimensions emerged prior to movement onset in active trials. Further validating these dimensions’ roles as a corollary discharge signal, we observed that their activity reversed in direction prior to the oppositely-directed return reach. For example, activity along the first example corollary-discharge dimension (**Figure 2B** top, the x-direction decoder) increased prior to the outward reach (i.e., dark blue trace, 0-degree, cosine = 1) and then reversed for the return (that is 180-degree, cosine = -1). Importantly, this demonstrated that the dimension was continuously present during movement, not simply at movement onset, suggesting it was a corollary discharge signal rather than motor preparation^36^.

The activity projected along these dimensions during passive trials (which were hidden from the decoders when identifying the signal) validated their consistency as a corollary discharge signal (**Figure 2B** right). Critically, activity along these dimensions reflected the voluntary return reach rather than the externally imposed perturbation. Returning to the example of the first dimension, during active trials, the dimension was positively modulated for outward, rightward reaches (dark blue trace, 0-degree) and then negatively modulated during the leftward return reach. During passive trials, the dimension during rightward bumps (dark blue trace) was not positively modulated, despite a rightward movement happening at that point—rather, it was negatively modulated corresponding to the upcoming voluntary leftward reach back to center. Further, activity remained near baseline prior to movement onset in passive trials, when there was no planned movement–another feature consistent with a corollary discharge signal. These patterns held for all reaching directions and dimensions (**Extended Data Figure 3**).

**Figure 3.**
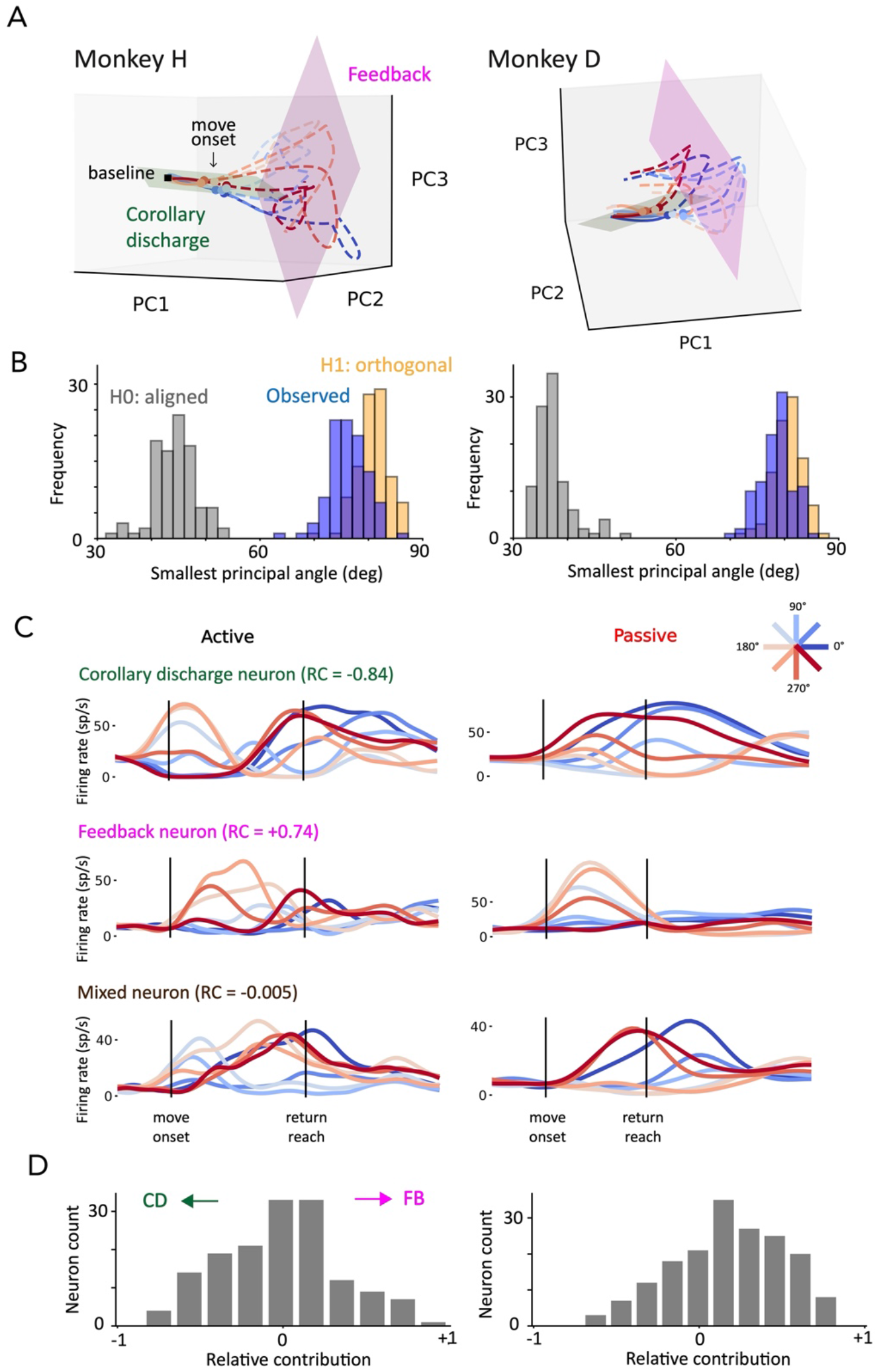
Corollary discharge and feedback signals are approximately orthogonal at the neural population level. **A**. Average neural trajectories for each reach direction during the first 500 ms of active trials in the three-dimensional space defined by top principal components. Corollary-discharge (green) and feedback (magenta) subspaces are overlaid as planes defined by their top two principal components within the same space. Prior to movement onset, neural activity occupied the corollary-discharge subspace. After, activity evolved within both signal subspaces. **B**. Distributions of the first principal angle across 100 Monte-Carlo trial splits. Shown are the angles between observed corollary-discharge and feedback subspaces (Observed, blue), and expected angles for orthogonal signals (Orthogonal, yellow) and identical underlying signals (Aligned, gray); see **Methods**. Observed distributions overlapped with those expected from orthogonal signals, with differences that were not significant (*p* = 0.28, 0.56 for Monkeys H, D, bootstrap test). **C**. Example single-neuron activity from Monkey H illustrating neurons that are corollary-discharge-like, feedback-like, or exhibit mixed responses. We list these neurons’ relative contribution (RC) indices, calculated based on their projections to the separate subspaces (see **Methods**). **D**. Distributions of the RC indices across the neural population.

### Orthogonal population subspaces support corollary discharge and feedback signals

#### Identifying feedback signals

To understand how sensory feedback is related to the corollary discharge signal we extracted, we next aimed to identify the feedback signal’s neural subspace. Since our extracted corollary discharge signal continuously led movement by ∼100 milliseconds, within the final 100 ms of a movement (when no future movement is occurring), we would expect minimal corollary discharge information. Thus, to find feedback signals unpolluted by corollary discharge signals, we aligned active trials to movement offset times and found the feedback signal’s constituent neural subspace within the 100-ms time window prior to movement offset (**Figure 2A**, magenta).

As we did with the corollary discharge signal, we verified the feedback signal’s identity by comparing its activity during active and passive trials (**Figure 2C**). As expected for a sensory feedback signal, activity along feedback dimensions emerged after movement onset in both trial types and reflected the movement independently of whether it was voluntarily or externally imposed. For example, in active trials, activity along the second example feedback dimension (**Figure 2C**; the y-direction decoder) increased first for the forward reach (i.e., light blue trace, 90-degree, sine = 1), then switched direction for the return (that is 270-degree, sine = -1). This dimension exhibited the same directional pattern in passive trials, even though the initial increase was now due a passive perturbation instead of a voluntary reach. Those results held for all reach directions and dimensions (total of 4 dimensions for Monkey H and 2 dimensions for Monkey D, **Extended Data Figure 4A**).

**Figure 4.**
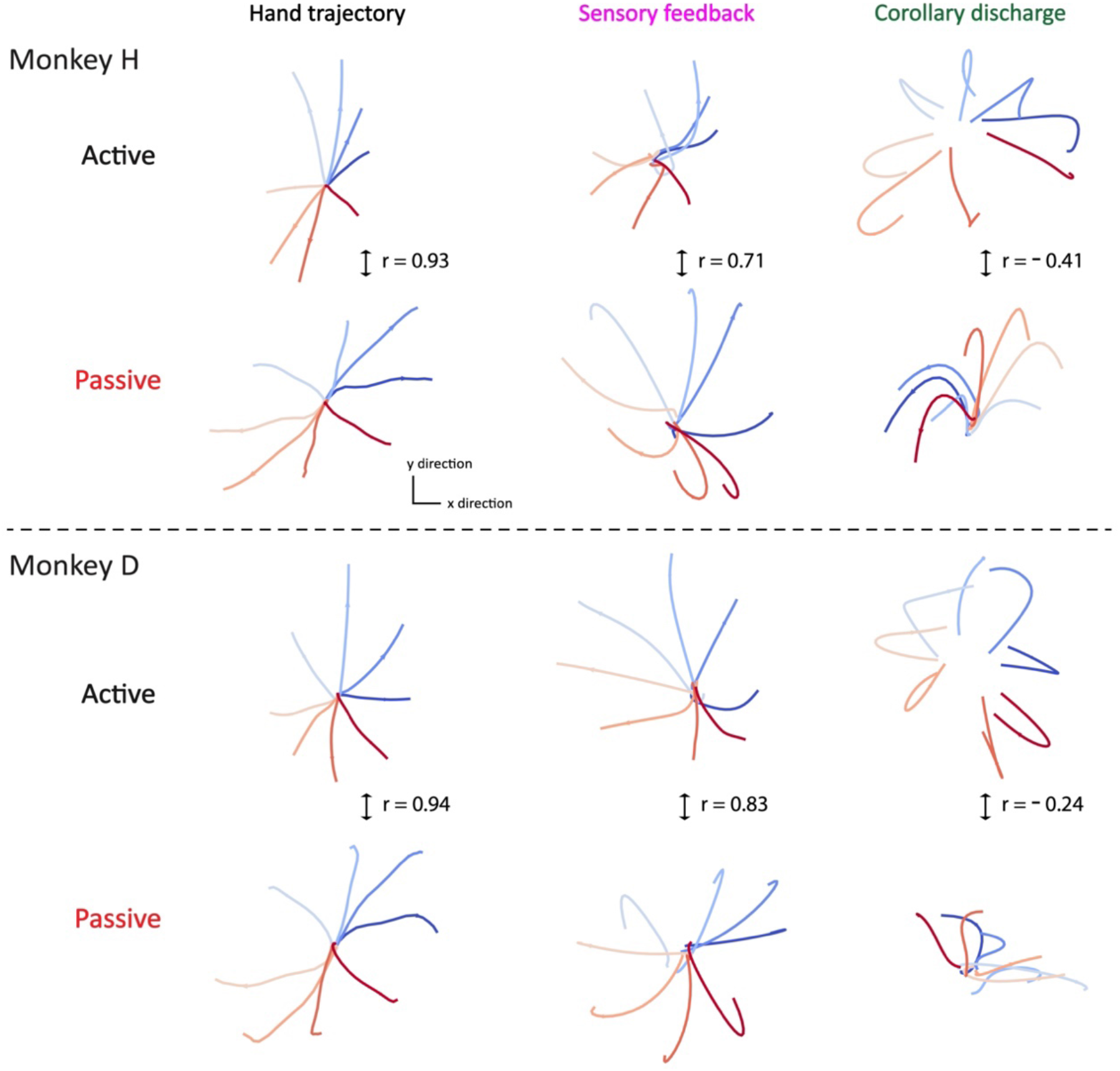
Feedback signals are similar across active and passive movements, whereas corollary discharge signals differ. Neural and kinematic trajectories during the time window [-100, 120] ms around movement onset of active and passive trials. Traces are averaged across trials within movement directions. Reported *r* values indicate Pearson correlation. *Left*: Hand trajectories (x-y velocity), which are similar between active and passive trials in this time window. *Middle*: Neural trajectories projected onto the top feedback dimensions corresponding to x- and y-direction decoders. Feedback trajectories were similar between active and passive conditions, as were the hand trajectories. *Right*: Neural trajectories projected onto the top corollary-discharge dimensions. In active trials, corollary-discharge trajectories led hand trajectories. In passive trials, they showed little alignment with hand trajectories and were weakly oppositely oriented relative to active trials, consistent with active trials reflecting reach-related activity and passive trials potentially reflecting return-related activity during the later part of the time window.

#### Orthogonal corollary discharge and feedback signals

After identifying the corollary-discharge and feedback signal’s subspaces, we aimed to discover how they related to one another. The large projection of neural activity into the corollary-discharge subspace before movement, with much less at the end of movement (and vice-versa for the feedback subspace) (**Figure 2B,C**) suggested that the subspaces are approximately orthogonal^36,37^. Indeed, visually inspecting their projections within a low-dimensional space of neural activity further suggests they are near orthogonal (**Figure 3A**).

We next quantified the extent to which these signals’ subspaces were orthogonal. To do so, we calculated principal angles between the corollary-discharge and feedback subspaces, comparing the observed angles to reference distributions constructed from different subsets of the same signal (representing well-aligned geometry) or from signals rendered orthogonal by projection (**Figure 3B**). The principal angle between corollary-discharge and feedback subspaces was substantially larger than would be expected from aligned signals (mean of 76 vs. aligned mean of 44 degrees for Monkey H, 78 vs. 37 degrees for Monkey D). In fact, it overlapped with what would be expected from two orthogonal signals (**Figure 3B**, blue vs. yellow distributions; **Extended Data Figure 6A** for additional monkeys/sessions).

#### Refining feedback signal estimation

Above, when seeking to test the relationship (i.e., the orthogonality) between corollary discharge and feedback signals, it was necessary to estimate the two signals independently. However, that approach has a strong limitation—neural activity toward the end of movement becomes sparse as muscle activation decreases and thus proprioceptive feedback weakens, which can compromise the estimation of the feedback signal. To address this, we exploited the approximate orthogonality between the two signals and adopted a stepwise extraction procedure. Specifically, we first found the corollary discharge signal as previously described, projected out its corresponding neural activity, and then estimated the feedback signal using the mid-movement period of active reaches, when feedback-related activity is more strongly elicited (**Figure 2D**). This procedure yielded a robust feedback subspace with six dimensions in both Monkeys H and D (**Extended Data Figure 4B**).

We confirmed that the feedback signal identified by this alternative method had the expected properties: as with the prior method, its directional modulation matched that observed during passive perturbation trials (**Figure 2E**), and its activity lagged kinematics in time. Importantly, the feedback subspace extracted using this stepwise procedure supported improved kinematic decoding compared to the original method (**Extended Data Figure 4C**), providing validation that it better captured feedback-related neural activity.

#### Single neurons make mixed contribution to corollary discharge and feedback signals

Two possible neural encoding mechanisms can give rise to the approximately orthogonal signal subspaces that we observed. One possibility is that the two signals might involve distinct subpopulations of neurons, such that single-neuron activity can be explained by one of the signals alone (“distinct encoding”). The other possibility is that the two signals might involve the entire neuronal population, with the majority of single neurons receiving a mixture of both signals^38^, organized onto orthogonal subspaces at population level (“mixed encoding”). We examined single-neuron activity and observed that while some neurons had predominantly corollary-discharge-like or feedback-like responses, many had features of both signals (**Figure 3C**). To differentiate the two mechanisms quantitatively, we characterized single neurons’ relative contributions to the two subspaces, defining the contribution index as the difference between each neuron’s contributions to feedback and corollary-discharge subspaces normalized by their sum. A contribution index of +1 indicates a feedback-only neuron and -1 indicates a corollary-discharge-only neuron^36^. In support of the mixed encoding mechanism, we found that the distribution of this index was centered near 0, indicating that many neurons contribute comparably to both corollary-discharge and feedback subspaces (**Figure 3D**; **Extended Data Figure 6B**).

### Corollary discharge explains neural activity differences between active and passive movements

Previous studies have shown that single neurons in somatosensory area 2 exhibit different activity patterns under active and passive movement contexts, even when the movement trajectories are comparable^19,27^. One plausible explanation is that the neural activity contains both corollary discharge and feedback signals during active movements, but only the feedback signal during passive movements, which results in the different neural activity patterns. However, this hypothesis has been challenging to test at the single-neuron level. Here, after dissociating the corollary discharge and feedback signals, we were able to compare their corresponding activity during the two trial types (**Figure 4**).

We examined the initial period when the active and passive hand kinematic trajectories were comparable (**Figure 4** left). As hypothesized, feedback signal trajectories were also similar (**Figure 4** middle, *r* = 0.71 and 0.83 for Monkeys H and D), both accurately reflecting hand trajectory at short latencies. On the other hand, corollary-discharge signal trajectories differed in the two types of trials (**Figure 4** right, *r* = -0.41, -0.24 for Monkeys H and D)—during active movements, corollary discharge signals led hand trajectories; during passive movements, corollary discharge signals contained little trajectory information. These findings supported the idea that corollary discharge contributes to the differences in neural activity patterns under active and passive movement contexts, while feedback signals are similar across these contexts.

### Signal integration compensates for feedback delay and improves body state estimation

To understand how the interaction between corollary discharge and feedback influences body state estimation, we compared movement kinematics decoding accuracy using neural activity within each signal’s subspace versus the combined subspace. We trained linear decoders to predict instantaneous hand velocity with a range of latencies relative to neural activity (**Figure 5A**). For the individual corollary discharge and feedback analyses, decoders were constrained to use non-negative weights on the extracted signal dimensions, preserving their original orientation. The combined decoder weights were unconstrained, allowing flexible mixing of corollary-discharge and feedback dimensions (see **Extended Data Figure 7** and **Methods**).

**Figure 5.**
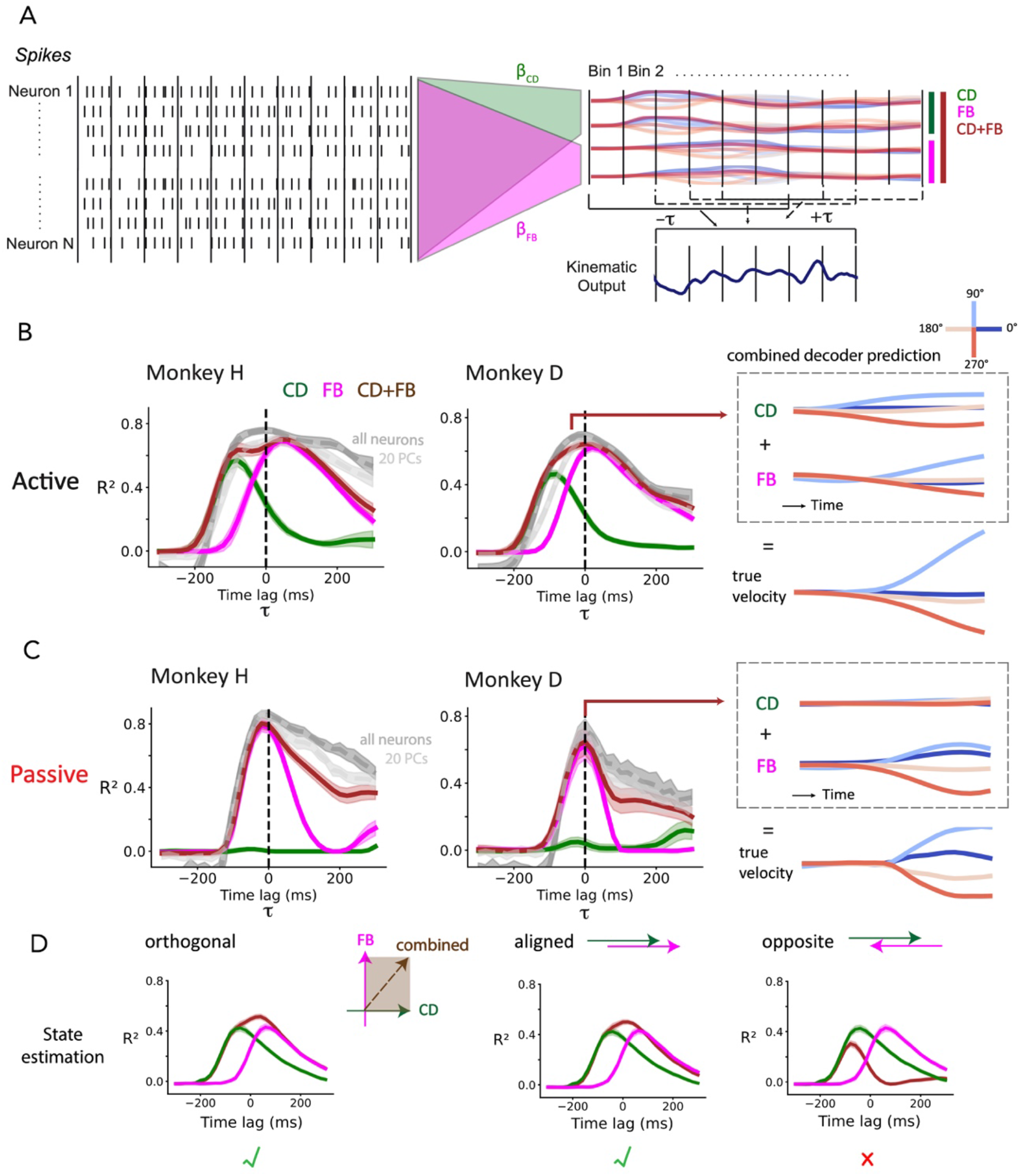
Signal integration improves body state estimation. **A**. Schematic of kinematic decoding analysis. We first extracted corollary-discharge and feedback subspaces from neural spiking activity using linear decoders (β_*CD*_and β_*FB*_), as in **Figure 2D**. Then, we used neural activity projected onto each signal’s subspace, or their joint subspace, to decode movement kinematics across different temporal lags between neural and kinematic variables. **B**. Velocity decoding in active trials. *Left*: Decoding performance as a function of neural-kinematic lag. Combined signals outperformed individual signals and approached the performance obtained using the full neural activity. Shading indicates standard deviation across 20 validation folds. *Right*: Example velocity predictions from the combined decoder for Monkey D, decomposed into contributions from corollary discharge and feedback signals, shown for 4 reach directions (out of 8) for clarity. The decoder trained at −40 ms lag is shown to illustrate the signals’ integration prior to peak feedback information. **C**. Same analysis as **B**, for passive trials, where integration has minimal benefit. *Right*: Example velocity predictions from the combined decoder for Monkey D, trained at 0 ms lag. **D**. Simulated decoding under different neural representational geometries. Corollary discharge and feedback signals were simulated with identical inputs but arranged in orthogonal, aligned, or opposite representations. Orthogonal and aligned geometries enabled successful integration, with combined signals outperforming individual signals and reproducing the decoding pattern in **B**. In contrast, opposite representations caused signal cancellation and poor decoding.

Our analyses centered on endpoint velocity, as it can be accurately decoded from area 2 neural activity, consistent with prior work showing that proprioceptor (muscle spindle) responses best relate to velocity^39^. Notably, our results also held for position and acceleration (**Extended Data Figure 8A,B**). Decoding accuracy was used as a proxy of the available kinematic information to be read out by downstream regions.

As expected, corollary discharge supported better decoding of future velocity, with optimal lags of -80 ms in both monkeys (**Figure 5B**), whereas feedback signals supported better decoding of past velocity (Monkey H: +40 ms, Monkey D: +20 ms). We note that, while these lags are reasonable, they should not be closely interpreted as physiological delays^40^ — still, the difference in optimal decoding lags between corollary discharge and feedback signals provides valuable information about their relative ordering. Interestingly, and in line with optimal control theories, combining corollary discharge and feedback signals enabled reliable kinematic estimation before the arrival of feedback (**Figure 5B**; brown curves reach high decoding R^2^ before magenta curves; **Extended Data Figure 9**). By probing the combined decoder’s use of corollary discharge and feedback signals within the decoder model, we found that both signals contributed to the predictions, indicating that the two signals were jointly used for state estimation (**Figure 5B** right, combined decoder prediction).

Notably, decoding accuracy using the combined subspace, despite its relatively low dimensionality (9-dimensional for Monkeys H and D), closely matched that obtained using the top 20 principal components, and even neared decoding accuracies of the full neural population (**Figure 5B**; brown curves resemble gray curves). These results suggest that integration of corollary discharge and feedback signals accounts for much of the kinematic information present in area 2 activity.

The two signals did not always operate in tandem. During passive perturbations, when there could be no predictable sensory outcome, the addition of corollary discharge did not offer more accurate or earlier decodability beyond that of feedback alone (**Figure 5C**; brown and magenta curves overlap). Consistent with this, the combined decoder used minimal corollary discharge signal to make predictions during passive trials, and kinematic predictions were made almost entirely by feedback (**Figure 5C**, combined decoder prediction).

Overall, corollary discharge provided earlier kinematic decodability, consistent with its role in helping to compensate for feedback delays. During active movements, integrating the two signals enhanced both decoding accuracy and the range of time lags over which kinematics could be reliably decoded, relative to either signal alone. This advantage disappeared during passive movements, when corollary discharge was absent.

How does neural population geometry impact this ability to integrate information effectively? To investigate, we simulated neural activity under three representational geometries —orthogonal, aligned, and opposite—and compared the resulting state estimation (**Figure 5D**). We modeled neural activity as the linear combination of past and future kinematics with Gaussian noise, using actual hand velocity from active trials, and fixed the signal lags by design (-80 ms for corollary discharge, +30 ms for feedback). Each signal alone could be used to decode velocity at its respective lag (**Figure 5D**, green and magenta curves), but their combination revealed key differences: both orthogonal and aligned signals improved decoding accuracy via integration and supported accurate decoding over an extended range of decoding lags, but opposite signals canceled and failed (**Figure 5D**, brown curves).

### Signal integration cancels sensory expectation and enhances perturbation detection

If both the aligned and orthogonal representational geometries are viable for accurate state estimation, why might the brain favor orthogonality? One potential answer lies in how the system responds to unexpected perturbations. While only the feedback signal reveals perturbations, the feedback signal’s concurrent representation of ongoing voluntary movement can mask the perturbation-related component and pose challenges for downstream regions to distinguish the two. Indeed, another important purpose of corollary discharge is the cancellation of predictable sensory feedback resulting from one’s own movements, as demonstrated by separate subfields of research^41–47^. We explored whether the approximately orthogonal corollary discharge and feedback signals could enhance perturbation detection via a cancellation-like mechanism.

To test this hypothesis, we recorded in monkey D during “reach-bump” trials, which included a brief assistive or resistive perturbation during the ongoing reach (**Figure 6A**). Perturbations were delivered in forward (**Figure 6B**, purple) or backward (yellow) directions, as the monkey reached either in the same or opposite direction. In this task, we would expect feedback activity to reflect a mixture of voluntary and unexpected (bump) components. Feedback alone would not differentiate bump direction well (purple vs. yellow traces), but the corollary discharge component could be used to cancel out the feedback related to the reach (**Figure 6B**). To generalize this example, we trained linear classifiers to separate opposite-direction bumps (e.g., forward vs. backward or left vs. right) across all reaching conditions (**Figure 6C**). The combination of corollary discharge and feedback information achieved higher accuracy than feedback signals alone.

**Figure 6.**
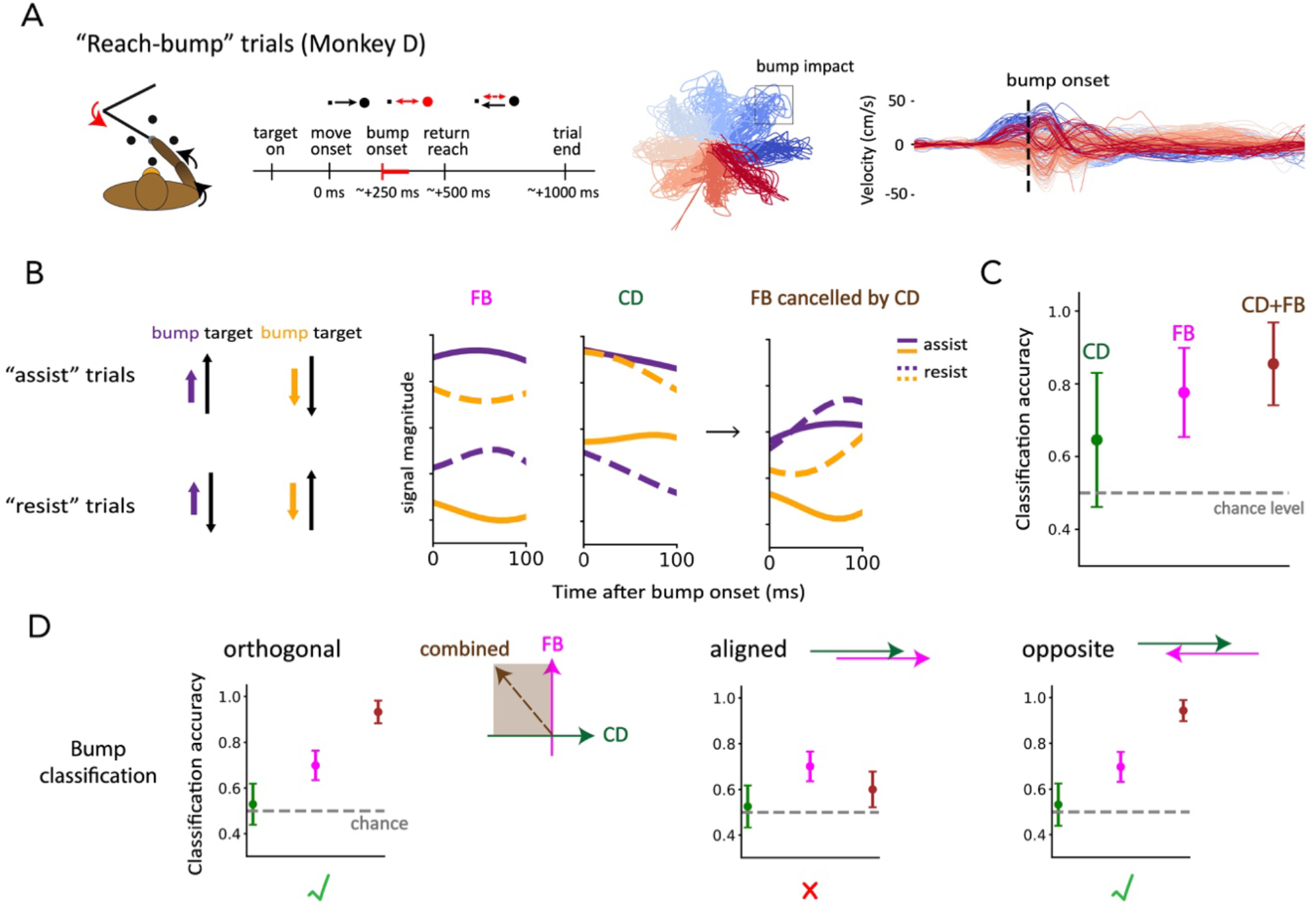
Signal integration enhances decoding of external perturbations. **A**. Perturbed reaching (“reach-bump”) task. *Left*: Trial structure. Black segments denote voluntary movements; red segments denote externally perturbed movements. *Middle*: Hand trajectories across individual trials, colored by reach direction. *Right*: Hand kinematics aligned to bump onset. **B**. Example illustrating how corollary discharge helps disambiguate perturbation direction. Neural activity is shown for y-direction dimensions of feedback and corollary discharge. Feedback alone does not differentiate bump directions (purple vs. yellow), and corollary discharge primarily reflects the voluntary reach direction. Subtracting corollary discharge from feedback reveals perturbation-related activity that separates the two bump directions. **C**. Perturbation classification in reach-bump trials. Incorporating corollary discharge improved classification performance relative to feedback alone. Mean classification accuracy was 0.65 ± 0.18 for corollary discharge, 0.78 ± 0.12 for feedback, and 0.86 ± 0.11 for the combined signals (mean ± SD across 100 validation folds). **D**. Simulated perturbation classification under different neural population geometries. Orthogonal and opposite geometries allowed combined signals to improve bump classification beyond feedback alone, matching experimental results in **C**, whereas aligned representations led to worse classification.

We again tested how different representational geometries (orthogonal, aligned, opposite) would affect perturbation decoding by adding simulated perturbations (transient Gaussian deviations) to the hand velocity from active trials (**Extended Data Figure 10**). Corollary discharge only reflected reaching-related velocity, whereas feedback also included perturbation-induced velocity. Combining the two signals improved bump classification beyond that of feedback alone when they were represented orthogonally or oppositely, matching our experimental results, while an aligned representation failed to improve bump classification (**Figure 6D**).

Together, we’ve shown that, while an aligned representation could allow integration for state estimation, and an opposite representation could allow integration for cancellation, only orthogonality enables both functions. Moreover, it was not simply that any multi-dimensional integration between corollary discharge and feedback (e.g. a 10-degree offset between the two) was sufficient to enable both accurate state estimation and cancellation—nearly-orthogonal relationships most robustly supported both functions (**Extended Data Figure 11A,B**). This flexibility makes orthogonality particularly advantageous for downstream decoders to flexibly extract different sources of movement information.

## Discussion

It has been long recognized that the integration of motor and sensory signals is crucial in supporting accurate perceptions^6,7,41^ and motor state estimation^10,11,14^, but the nature of their interactions within cortical populations has remained unclear. In this study, we investigated how corollary discharge and feedback signals interact in somatosensory area 2 by analyzing neural recordings from monkeys performing active and passive center-out reaching tasks. By considering their distinct onset times, we were able to dissociate corollary discharge and feedback signals and found that they occupy approximately orthogonal neural subspaces, resulting in their demixed representations during movement. In line with optimal feedback control theories, the integration of the two signals allowed earlier accurate kinematic estimates during active movements. To our knowledge, this is the first study to provide neural evidence, based on spiking activity, that the sensory cortex integrates corollary discharge and sensory feedback to support state estimation.

Furthermore, integrating corollary discharge and sensory feedback helped to discriminate external inputs by cancelling self-generated sensory reafference, and our simulation results showed that orthogonality underlies flexible signal usage across both kinematic estimation and perturbation detection. That is, orthogonality preserves distinct sensorimotor representations while allowing their selective combination when needed. This suggests that orthogonality may be a core cortical strategy for supporting flexible integration without loss of signal specificity – synthesizing distinct theories on the role of corollary discharge in optimal feedback control^10–12^ and the suppression of predictable sensory input^42–45^, which have previously been considered in isolation.

Recent studies have likewise revealed other orthogonal latent subspaces in brain activity. The hypothesized functional advantages of such orthogonality include the separation of different computation phases^36,37,48^, concurrent representation of multiple information modalities^49–52^, and flexible routing of information^53^. Our findings thus align with a growing body of evidence suggesting that orthogonality is a widespread principle governing distinct neural representations across the brain.

### Differences across monkeys

We observed variability in the strength of corollary discharge signals across animals: signals were most pronounced in Monkeys H and D, and we therefore focused our downstream analyses on these monkeys to characterize signal integration. Nonetheless, we continued to observe consistent qualitative results in Monkeys C and L (**Extended Data Figures 6, 9**), including the approximate orthogonality between corollary discharge and feedback signals and modest benefits of their integration, consistent with their limited corollary discharge signals to start with. Multiple factors may have contributed to this difference. First, recordings from Monkeys C and L included fewer neurons (57 and 88 versus 153 and 176 in Monkeys H and D), likely contributing to their lower kinematic decoding accuracy at the raw population level, prior to any signal extraction (**Extended Data Figure 9**). Relatedly, they may have had fewer responsive neurons and different response properties^19^. Finally, Monkeys C and L performed a task with only four reach directions rather than eight, which may have provided less directional structure in behavior from which to extract corollary discharge-related activity. Together, these factors likely account for the quantitative differences observed across animals, while our main conclusions generalize qualitatively across datasets.

### Sources and interpretations of the extracted signals

While we interpret the extracted signals as reflecting corollary discharge and sensory feedback, we cannot precisely know their sources, and alternative interpretations are possible. The ‘corollary discharge’ signal is consistent with input from motor cortex (M1) given its timing and directional specificity, although it could arise from another motor region. While our corollary discharge signal shares some features with purely ‘preparatory’ signals^36^, such as its direction specificity prior to movement onset, our signal’s persistence throughout movement makes this alternative explanation unlikely. And while global arousal-related activity^31–33^ may occur prior to and during movements, we would not expect this to have the direction-specific structure observed in our corollary discharge signals.

While we view the ‘feedback’ signal as predominantly reflecting peripheral sensory input, it may also include other components. When feedback was estimated using neural activity aligned to movement offset, the estimation window may have included some corollary discharge related to braking. However, this contribution would be limited to a small fraction of the window and therefore weakly expressed in the extracted signal. More generally, any mix of corollary discharge with feedback would cause the angle between the two signals to be underestimated, suggesting that, if anything, the signals are more orthogonal than we reported.

### Interareal communications of the integrated signals

Our findings demonstrate that area 2 contains complementary corollary discharge and sensory feedback signals in a form suitable for integration, rather than directly establishing that integration is performed there. Although we cannot determine whether the brain explicitly combines these signals along the specific dimensions we identified, their joint availability enables multiple downstream computations. We hypothesize that the combination of feedback and corollary discharge signals—encoding current body state estimates—may be routed to higher-order parietal and (indirectly) to prefrontal regions involved in learning and internal model updating^54–56^, whereas the comparison between the two signals—potentially encoding sensory prediction errors—may be transmitted to motor areas to additionally support online motor control^57–61^. This is consistent with the anatomical evidence that area 2 projects to both motor and parietal areas, making it an ideal site for rapid online feedback control and error monitoring^24,25^.

Given that we recorded only from area 2, we cannot directly assess these hypotheses. With the growing prevalence of multi-area recordings and perturbation approaches, future studies could more directly evaluate the integration and utilization of these signals across multiple cortical networks^31,32^.

## Author Contributions

X.A. and J.I.G performed analyses. R.H.C. and K.P.B. collected the data. R.H.C., K.P.B., and L.E.M designed the experiments. X.A. and J.I.G. wrote the manuscript. All authors contributed to manuscript editing.

## Acknowledgements

We thank Andrew Miri for helpful comments and discussions. This work was supported by NIH ROONS119787 (X.A, J.I.G.), F32MH120893 (K.P.B.) and NIH R01NS095251 (K.P.B, R.H.C, L.E.M). This research was also supported in part by grants from the NSF (DMS-2235451) and Simons Foundation (MPS-NITMB-00005320) to the NSF-Simons National Institute for Theory and Mathematics in Biology (NITMB).

## Declaration of Interests

The authors declare no competing interests.

## Methods

All surgical and experimental procedures were fully consistent with the guide for the care and use of laboratory animals and approved by the institutional animal care and use committee of Northwestern University under protocol #IS00000367.

### Behavioral tasks

We recorded from four rhesus macaques (Monkey H, D, C, and L) performing a planar center-out reaching task using a two-link manipulandum (protocol originally described in ^27^; datasets for Monkey H, C, and L first published in ^19^). The animal controlled a handle within a 20×20 cm workspace to acquire visually displayed targets, and a pulse of juice or water was rewarded after each successful reaching trial.

Monkeys H and D performed reaches to eight peripheral targets, uniformly spaced at 45° intervals (0°–315°), for which we presented results in the main figures. Monkey C and L performed reaches to four peripheral targets, spaced at 90° intervals. In all animals, trials began with the monkey holding a central target for a variable duration. Trials were either active reaching (“active”), passive perturbation (“passive”), or perturbed reaching (“reach-bump”) trials.

Active and passive trials followed the previously published protocol and were randomly interleaved during a session. Briefly, in active trials, the monkey reached from a central target to one of the peripheral targets and then returned to the center. In passive trials, a brief mechanical perturbation was delivered during the center-hold period in one of the target directions; the monkey actively returned to the center afterwards. The mechanical perturbation in passive trials was 2 N for Monkey H, C, L and 1.5 N for Monkey D, with force magnitudes chosen to produce movements which matched the monkey’s hand kinematics during the 120 ms after movement onset in active trials.

Reach-bump trials were collected only in Monkey D and were also randomly interleaved with active and passive trials. These trials combined elements of both the active and passive trial types. The monkey first initiated an active reach toward a peripheral target, as in an active trial. Then, a brief mechanical perturbation of the same strength as passive-trial perturbations was delivered during the middle of the reach, 250 ms after go cue. Perturbations included in the analyses were either assistive (directed toward the target) or resistive (directed opposite the target direction). Perturbations orthogonal to the target direction were also present but were excluded from analyses, as we were specifically concerned with cancellation of self-generated sensory consequences, and perturbations aligned with the movement axis were most directly relevant. After the perturbation, the monkey continued reaching to the target and returned as in a normal active trial.

A trial was successful once the monkey reached the outer target. Only successful trials were included in analyses. We recorded the joint angles of the manipulandum and calculated the animal’s endpoint kinematics based on them.

### Data processing

#### Neural data acquisition and pre-processing

We implanted Utah arrays (Blackrock Neurotech) in the arm representation of area 2 of somatosensory cortex of each monkey, as detailed in ^19^. We recorded 96 channels at 30kHz, using a Cerebus recording system. Neural spikes were detected online using threshold crossing and manually sorted using Plexon offline sorter into putative units based on waveform features and inter-spike interval criteria. Neurons with firing rate below 0.1 Hz were excluded from analyses.

We used tools from Neural Latents Benchmark^62^ for dataset loading and formatting. For all analyses, neuron spikes were binned using 10-ms bin size. In **Kinematic decoding**, we used a 40-ms Gaussian filter to smooth neural activity. Otherwise, we used unsmoothed neural activity in **Corollary discharge and feedback signal extraction** and **Perturbation classification** to avoid temporal leakage due to filtering.

#### Behavioral alignment

We used single-trial data for all the analyses. We estimated movement onset by first locating the first peak in hand acceleration occurring starting after the go cue for active trials (or after 50 ms pre-bump delivery time for passive trials), then stepping backward in time to the first point where acceleration fell below 20% of the peak value. If no valid acceleration peak was found, onset was instead defined as the first time that hand speed exceeded 5 cm/s.

We similarly estimated movement offset by locating the last acceleration peak within the movement period and then stepping forward in time to the first point where acceleration dropped below 20% of that peak. If no valid peak was detected, offset was defined as the first time that speed fell below 5 cm/s.

### Neural analysis

#### Sliding-window t-test for onset detection

We determined neural activity onset (**Figure 1C**) by performing a t-test across trials comparing the mean firing rate in each 20-ms moving window (10-ms step size) to a baseline window (−250 to −150 ms relative to movement onset). For each trial, spike activity was first averaged across neurons to obtain a single population firing-rate time series. We performed the moving-window comparisons over the −150 to 0 ms interval preceding movement onset. Onset time was defined as the earliest time bin from which all subsequent bins were significantly different from baseline (paired two-tailed t-test, p < 0.05). This consecutive-significance criterion was used to reduce spurious detections arising from fluctuations in firing rate.

#### Corollary discharge and feedback signal extraction

We identified low-dimensional neural subspaces associated with specific signals (corollary discharge and feedback) by fitting linear ridge-regression models that mapped population activity in particular time windows to movement direction. This procedure was applied separately to the sine and cosine components of reach direction.

Let *X* ∈ ℝ^*Tr×N*^ denote the unsmoothed neural activity averaged within a designated time window and then z-scored across trials (with *Tr* trials and *N* units). Let each element of *Y* ∈ ℝ^*Tr*^ denote a scalar directional signal, defined as either the sine or cosine of the reach direction. To estimate a signal axis, we fit a linear model

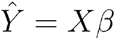

where *β* ∈ ℝ^*N*^ was estimated by minimizing a ridge-regularized least-squares objective,

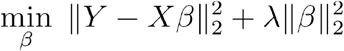

Predictive performance for each axis *β* was quantified using 10-fold cross-validation stratified by movement direction. For each fold, the regularization coefficient *λ* was selected by a logarithmic grid search within the training set with 5-fold cross-validation.

After fitting an axis *β*, we removed the neural activity in the corresponding subspace by projecting the population activity*X* onto the orthogonal complement of the subspace spanned by *β*. This was accomplished using the projection matrix

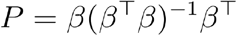

and the residual activity was computed as

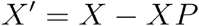

The residual activity *X*′ was then used to iteratively refit additional axes, continuing until the cross-validated coefficient of determination (R^2^) for the next axis dropped below zero. This procedure was performed separately for the sine and cosine components, yielding a set of axes that together spanned the subspace associated with the signal of interest.

We applied this framework to extract subspaces corresponding to corollary discharge and feedback signals. The corollary discharge signal was extracted using the neural activity in the 100-ms window preceding movement onset. When testing orthogonality of the feedback and corollary discharge signals (**Figure 3B**), we extracted the feedback signal using the activity in the 100-ms window preceding movement offset. After establishing orthogonality, in order to find a more robust feedback signal, the feedback signal was extracted using neural activity 200–400 ms after movement onset; in this case, the activity was first projected onto the orthogonal complement (nullspace) of the corollary-discharge subspace to isolate feedback-related variance.

#### Quantifying the relationship between corollary discharge and feedback signals

We quantified the geometric relationship between the corollary-discharge and feedback subspaces using principal angles, which quantify the best possible alignments between two subspaces embedded in a higher-dimensional neural activity space. Principal angles are defined sequentially as the angles between pairs of unit vectors (one from each subspace) chosen to minimize the angle between them while remaining orthogonal to previously selected pairs. By construction, this method yields an ordered sequence

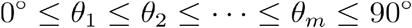

where *m* is the smaller of the two subspace dimensions, and *θ*_1_ is the first (smallest) principal angle. In our analyses, we focused on this smallest angle, which provides the most sensitive measure of subspace alignment: small values indicate substantial overlap, whereas values closer to 90° are more orthogonal.

To assess whether corollary discharge and feedback signals occupy aligned or distinct neural subspaces, we compared the distribution of their first principal angles to two reference distributions representing the cases of identical and orthogonal underlying signals. We did this with a Monte Carlo resampling procedure consisting of 100 random 50/50 splits of the trial set, stratified by movement direction. On each iteration, we partitioned trials into two complementary halves (“split 1” and “split 2”). We fit corollary-discharge and feedback subspaces separately within each half, denoting the resulting estimates as CD_1_ and FB_1_ for split 1, and CD_2_ and FB_2_ for split 2.

We then formed three distributions of first principal angles:

1. Observed (CD_1_ vs. FB_1_) – corollary-discharge and feedback subspaces estimated independently from the same split, to quantify their direct alignment;
2. Aligned (CD_1_ vs. CD_2_) – corollary-discharge subspaces estimated independently from two complementary splits, to establish the expected distribution when recovering the same underlying signal;
3. Orthogonal (CD_1_ vs. FB_2_, where FB_2_⊥CD_2_) – corollary-discharge subspace from split 1 compared to the feedback subspace refit on split 2 after removing components aligned with the corollary-discharge subspace (yielding FB_2_⊥CD_2_), to establish the expected distribution when underlying signals are orthogonal.

We assessed the significance of differences between angle distributions using a bootstrap test. We randomly sampled one value from each angle distribution and computed the difference 1,000 times. A two-sided p-value was estimated as twice the smaller of the proportions with positive or negative differences.

#### Relative contribution index

We quantified each neuron’s contributions to the two signals using a relative contribution index (**Figure 3C,D**), defined as

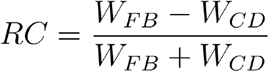

where *W*_*CD*_ and *W*_*FB*_ are the neuron’s average absolute decoder coefficient across corollary-discharge and feedback dimensions, respectively.

#### Kinematic decoding

To quantify how specific signals predicted hand kinematics, we trained linear ridge-regression decoders to map neural activity to instantaneous hand velocity. Let

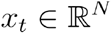

denote the vector of neural activity from *N* units (or signal dimensions) at time *t*, and let

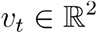

denote the corresponding planar hand velocity (horizontal and vertical components). Velocity decoding was performed over a fixed window from -100 to +120 ms relative to movement onset for both active and passive trials, where the hand kinematics were comparable by experimental design. Neural activity was shifted relative to this fixed kinematic window by a temporal lag *τ*, pairing velocity at time *t* with neural activity at time *t* + *τ*. Lags were sampled from −300 to +300 ms in 20-ms increments.

For each lag, neural activity was z-scored using training-set statistics, and hand velocity was mean-subtracted based on training trials. Timepoints were concatenated across trials to form

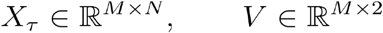

where *M* is the total timepoints across trials. We fit Ridge-regularized decoders of the form

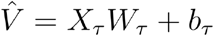

by minimizing

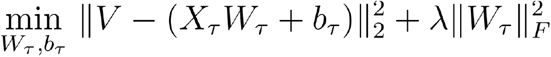

At each lag, five decoders were trained using:

1. the corollary-discharge (CD) subspace,
2. the feedback (FB) subspace,
3. the combined (CD+FB) subspaces,
4. the full neural population, and
5. the top 20 principal components (PCA fit to all timepoints within a session).

Because corollary-discharge and feedback axes were defined by regression onto the sine and cosine of the reach direction, their decoders (1) and (2) were sign-constrained to preserve the directional meaning of the signal. That is, decoder weights in were restricted to be non-negative to prevent arbitrary sign inversions and maintain a consistent mapping between signal activity and kinematics. Combined signal, population, and PCA decoders were unconstrained.

Decoder performance at each lag was evaluated using Monte Carlo cross-validation with 20 90/10 train-test splits at the trial level and stratified by movement direction. For each split, the regularization parameter was selected by logarithmic grid search within the training set with 5-fold cross-validation. Repeating this for all lag values yielded the R^2^s as a function of lag.

#### Perturbation classification

To quantify how corollary discharge and feedback signals related to externally applied perturbations, we trained logistic-regression classifiers to decode bump direction from neural activity in reach-bump trials. Let

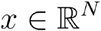

denote the neural activity from *N* signal dimensions averaged over the 50-ms window following bump onset (25–75 ms), and let

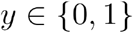

denote the binary perturbation-direction label (e.g., forward vs. backward). We analyzed the 25– 75 ms time window to capture early cortical processing of the perturbation while minimizing contributions from later voluntary responses (>100 ms)^63^. During this window, the cortical circuits would be expected to contain both reach-related motor signals and perturbation-related sensory input, and are thus well suited to test whether corollary discharge helps differentiate perturbation during an ongoing reach.

The classification analysis was performed separately for each direction axis, defined as a target direction and its 180° opposite (0°/180°, 45°/225°, 90°/270°, and 135°/315°), containing both assistive and resistive bumps.

Within each cross-validation split, features were z-scored using the training-set mean and standard deviation. For each direction and signal space, this yielded

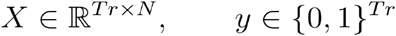

where *Tr* is the number of trials. We fit classifiers of the form

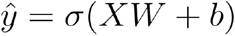

where *σ* (·) is the logistic function. The parameters *W* and *b* were estimated by minimizing the logistic-regression loss

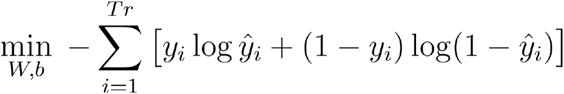

Classification was performed using:

1. the corollary-discharge (CD) subspace,
2. the feedback (FB) subspace,
3. the combined (CD+FB) subspaces.

No sign constraints were applied to the perturbation classifiers.

Classifier performance was evaluated using 100 Monte Carlo cross-validation folds with 50/50 train-test split. Splits were stratified by the four reach-bump categories (opposite target directions × assist/resist bumps) to ensure balanced sampling of behavioral conditions. Accuracy was computed on the held-out data and averaged across the direction axes.

#### Simulations

##### Simulation for kinematic decoding

To examine how the geometry between corollary-discharge and feedback subspaces affects kinematic decoding, we constructed a model of neural activity driven by recorded hand velocity from Monkey D during the same fixed window as in **Kinematic decoding**, from -100 to +120 ms relative to movement onset in active trials.

A population of 100 simulated neurons were assigned separate corollary-discharge and feedback weight matrices, *W*_CD,_ *W*_FB_ ∈ ℝ^100×2^. To generate these weights, a 100 × 4 matrix was sampled from a standard normal distribution and orthonormalized; the first two columns defined the corollary-discharge weight matrix *W*_CD_ and the remaining two columns defined an orthogonal complement *W* _⊥_. Each column corresponded to x- or y-components of velocity. Alternative geometries were generated by constructing feedback weights as a linear combination of the corollary-discharge subspace and its orthogonal complement according to

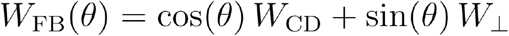

where *θ* controlled the relative alignment between corollary-discharge and feedback representations. This construction yielded aligned (*θ* =0°), orthogonal (*θ* =90°), opposite (*θ* =180°), and intermediate partially aligned geometries.

Neural activity was simulated by linearly projecting hand velocity signals into the full neural activity space by multiplying velocity with the CD and FB weight matrices. We added independent Gaussian noise of fixed magnitude (std = 10 cm/s) to the CD and FB streams of velocity inputs prior to projection, to model distinct noise sources associated with corollary discharge and sensory feedback pathways. After projection into neural space, additional independent Gaussian noise was added to the CD and FB activities to model neural noises. We varied the standard deviations of neural noises to assess the robustness to noise under different geometries (**Extended Data Figure 11A**).

We designed the simulation such that the signal lags match their empirical peak decoding times (**Figure 5B**): CD activity was driven by hand velocity at a −80 ms lag and FB activity by velocity at a +30 ms lag. For each lag, the segment of simulated neural activity corresponding to the decoding window was extracted and decoded using the same ridge-regression procedure and cross-validation described in **Kinematic decoding**. R^2^-versus-lag curves were computed separately for CD-only, FB-only, and combined activity.

#### Simulation for perturbation classification

We used a similar model framework to test how CD–FB geometry affects perturbation decoding. Recorded hand velocity during active trials of Monkey D again served as the baseline input. Synthetic perturbations were introduced by adding a Gaussian velocity pulse (40-ms standard deviation, fixed peak magnitude of 20 cm/s) centered at T = 325 ms after movement onset, in the assistive or resistive direction relative to the ongoing reach. With this parametrization, the effective onset of the perturbation occurred approximately two Gaussian widths earlier at T ≈ 245 ms, which yielded simulated velocity profiles that closely matched those observed in experimentally recorded reach-bump trials (**Extended Data Figure 10**).

The neural population construction according to different CD–FB geometries followed the same framework as in Simulation for kinematic decoding (**Extended Data Figure 11B**). CD activity was driven by the unperturbed velocity at a −80 ms lag, whereas FB activity was driven by the perturbed velocity at a 0 ms lag–its peak decoding latency during passive movements (**Figure 5C**). As described in **Perturbation classification**, for each trial, neural activity was averaged over 25– 75-ms window following perturbation onset (T = 270–320 ms) to form the feature vector. Perturbation direction (assistive or resistive) was classified using the logistic-regression framework, applied separately to CD-only, FB-only, and combined activity.

## Extended Data

**Figure 1.**
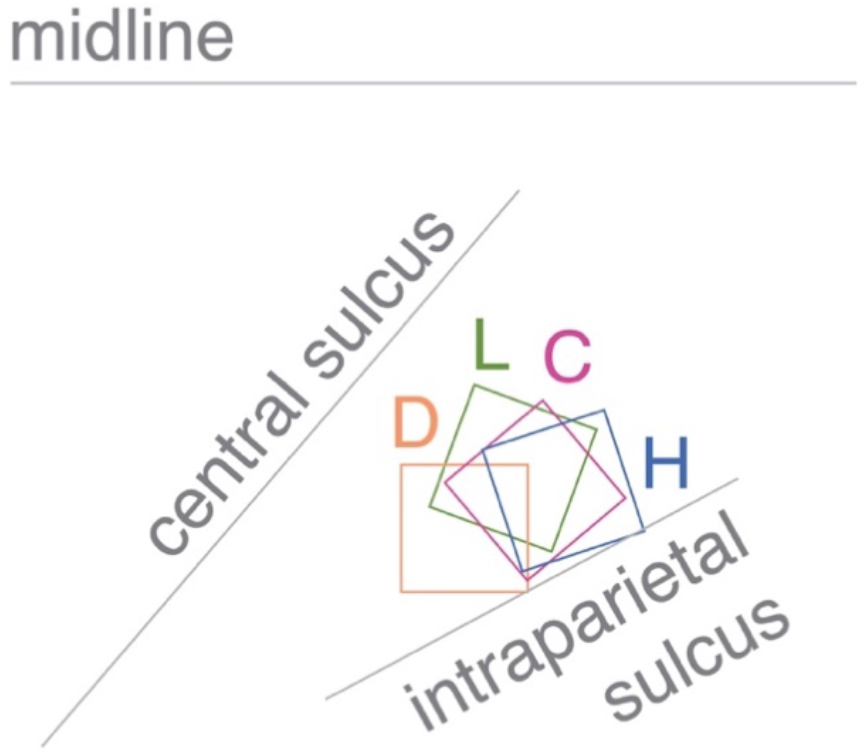
Locations of the chronic implants in somatosensory area 2 of four monkeys.

**Figure 2.**
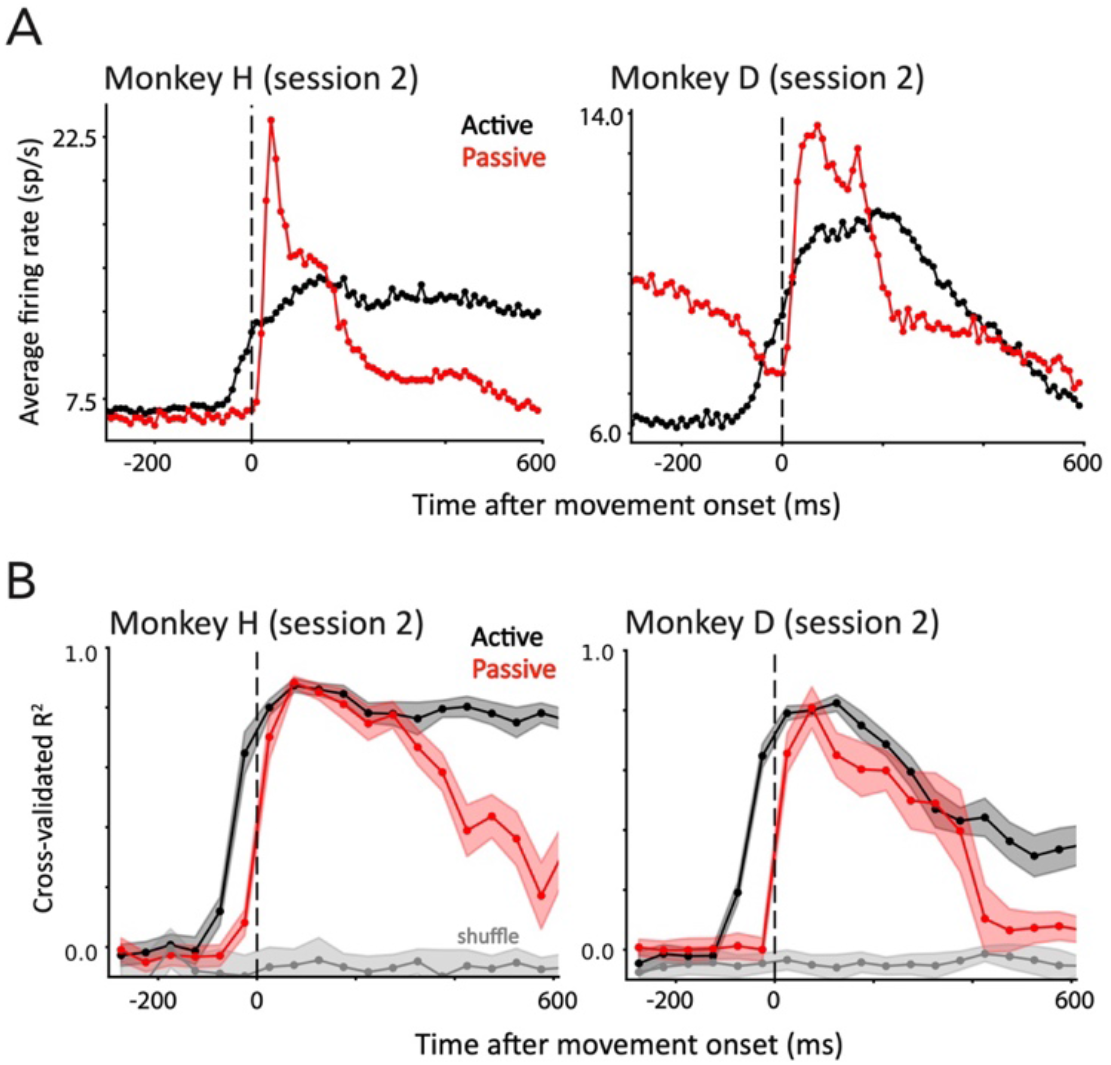
Evidence of corollary discharge from second sessions of Monkey H and D. Same format as Figure 1. **A**. Average neural firing rates during active and passive trials. Activity onset time in active trials was -70 ms for the second session of Monkey H, and -80 ms for Monkey D. **B**. Decoding accuracy at each time bin. Shading shows standard deviation.

**Figure 3.**
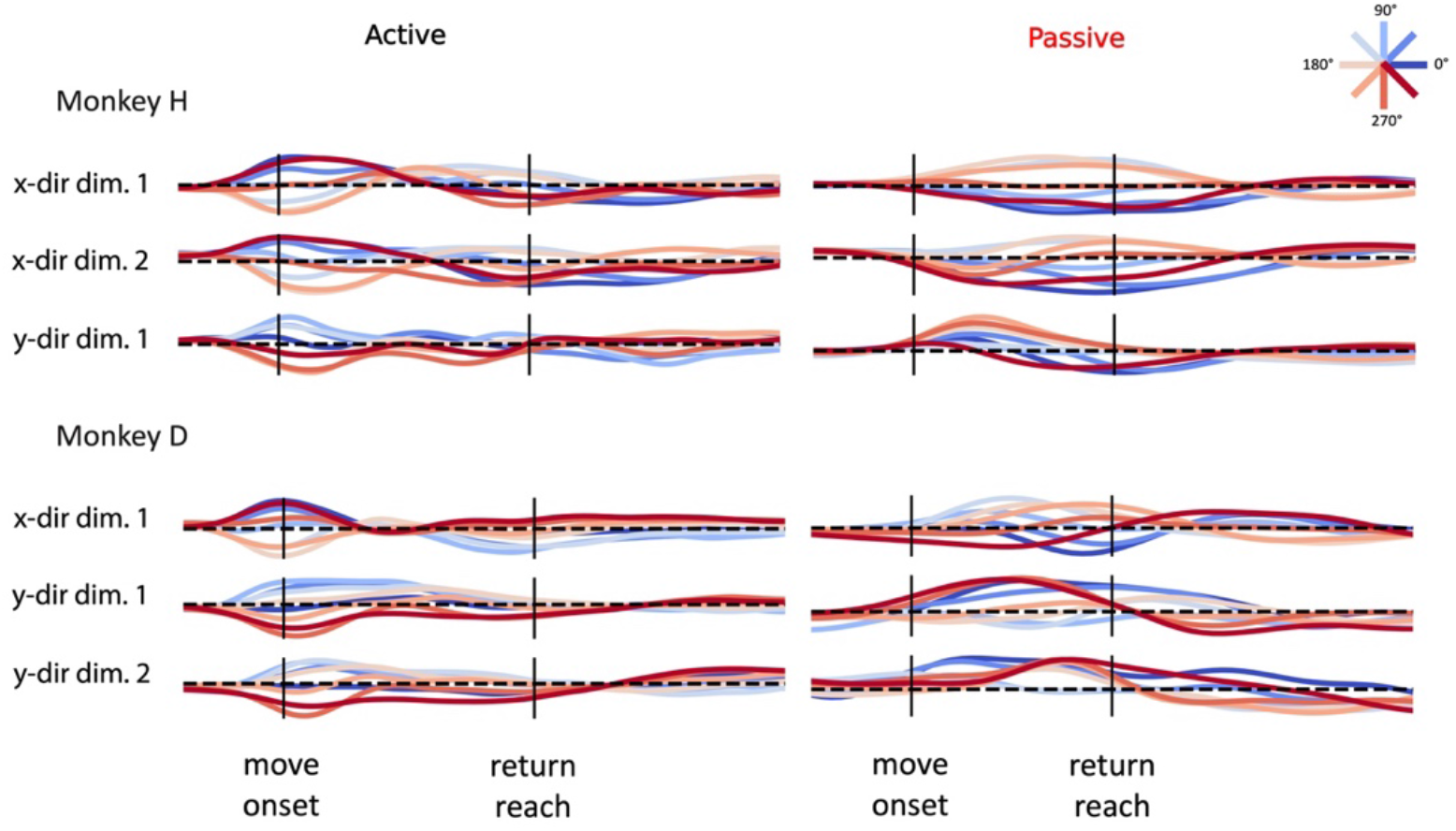
**All corollary-discharge dimensions identified for Monkey H and D**, expanding on those shown in **Figure 2B**. Neural activity projected onto each identified corollary-discharge dimension during active and passive trials. Dimensions are ordered by their order of extraction, i.e., x-dir dim. 1 corresponds to the first decoder fit to predict horizontal reach directions, dim. 2 corresponds to the second decoder after projecting out the activity related to the first dimension. Later dimensions have lower decoding accuracy and less directional information. We repeated this extraction procedure until the validation accuracy of the next dimension dropped below zero.

**Figure 4.**
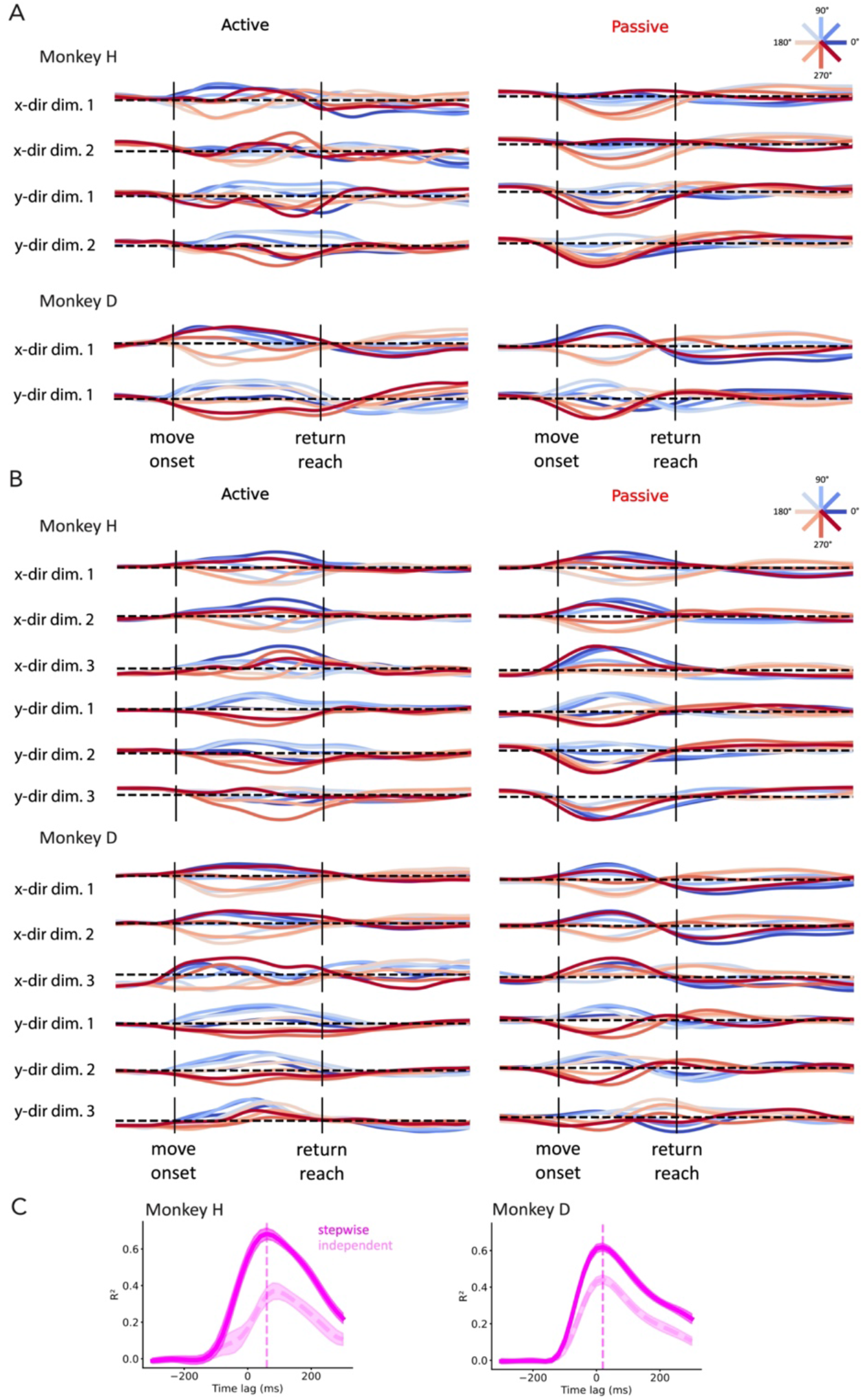
Comparison of feedback dimensions extracted using independent and stepwise procedures. **A**. All feedback dimensions identified with the independent extraction procedure for Monkeys H and D, expanding on those shown in **Figure 2C**. Same format as **Extended Data Figure 3. B**. All feedback dimensions identified with stepwise extraction procedure for Monkey H and D, expanding on those shown in **Figure 2E. C**. Comparison of velocity decoding in active trials between the feedback dimensions extracted in independent and stepwise manners. Decoding performance was plotted as a function of neural-kinematic lag (as in **Figure 5B**), where a positive lag means the neural signal lags kinematics. Both procedures captured similar, positive peak decoding lags. Feedback dimensions extracted with stepwise procedure (solid magenta) achieved higher decoding accuracy.

**Figure 5.**
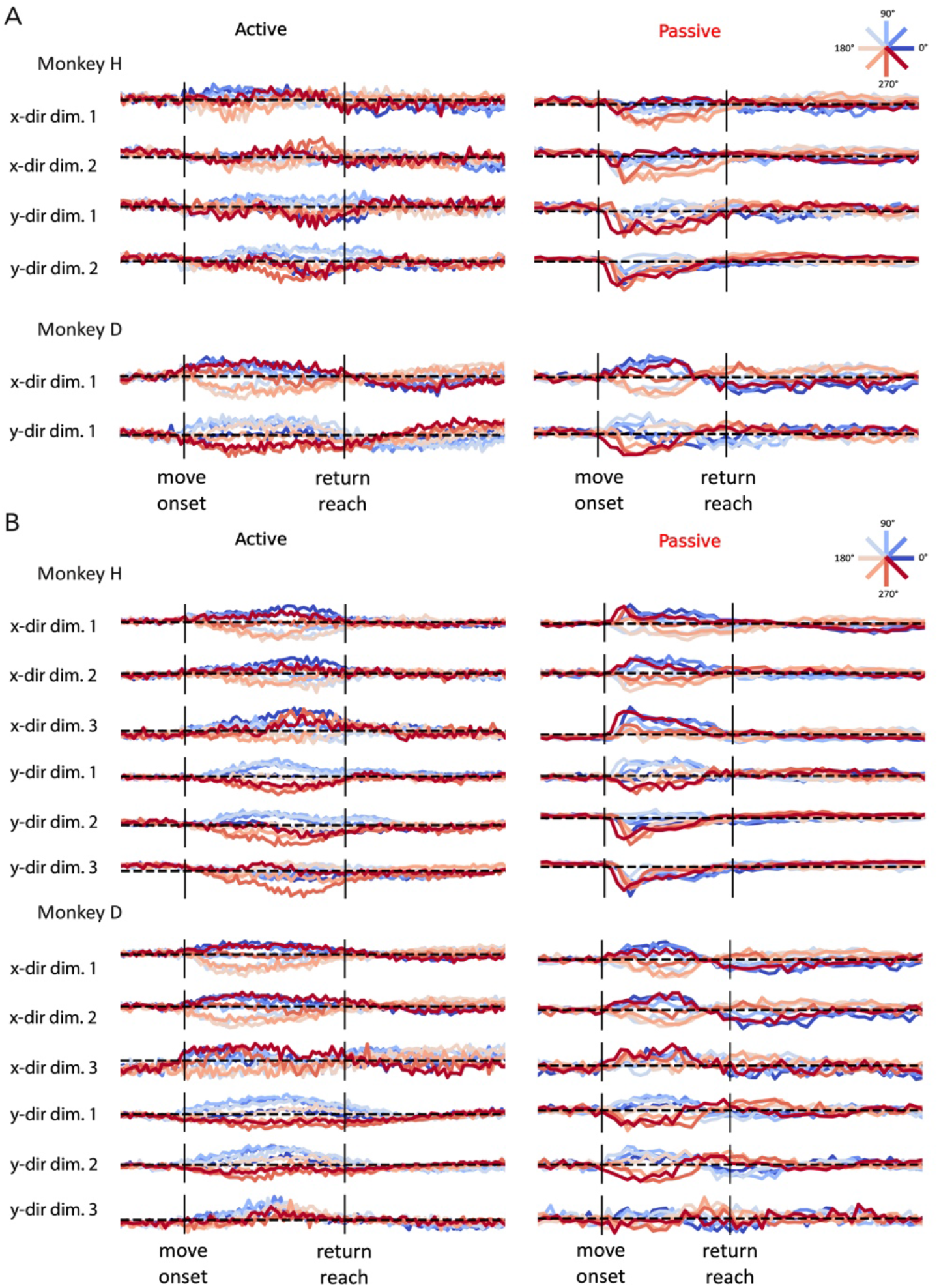
Unsmoothed traces of feedback dimensions extracted using independent and stepwise procedures. Same format as Extended Data Figure 4 A,B. **A**. All feedback dimensions identified with the independent extraction procedure for Monkeys H and D. **B**. All feedback dimensions identified with stepwise extraction procedure for Monkey H and D.

**Figure 6.**
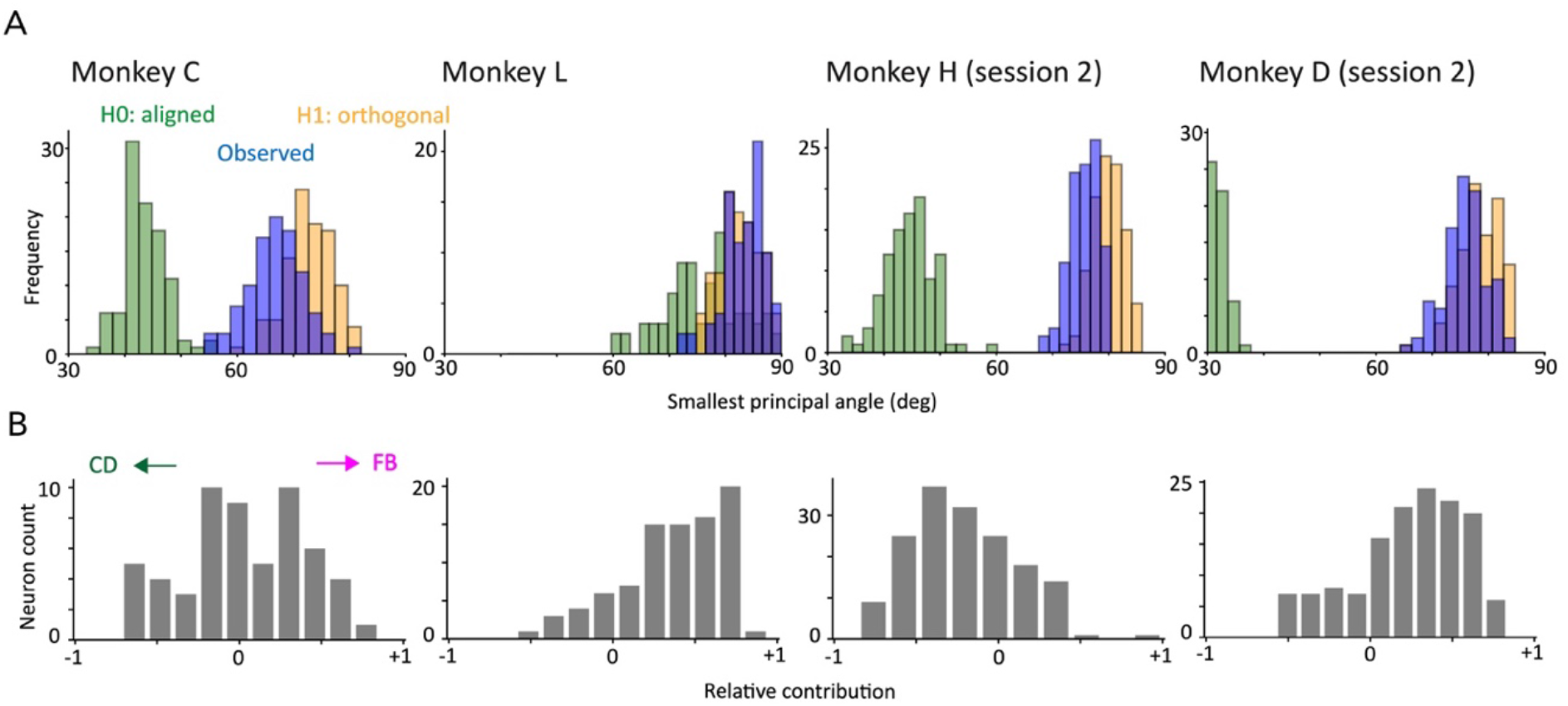
Quantifying relationship between corollary discharge and feedback signals in Monkeys C and L and second sessions of Monkey H and D. Same format as Figure 3. **A**. Distributions of the first principal angle across 100 Monte-Carlo trial splits. **B**. Distributions of the relative contribution indices across the neural population. Note: Corollary discharge could not be reliably extracted in Monkey L (failed in 21 of 100 trial splits), indicating a weak signal. Accordingly, in panel A the distribution for identical signals (green; between corollary discharge signals identified in two halves of trials) overlaps with that for orthogonal signals (blue), and in B most neurons appear feedback-driven.

**Figure 7.**
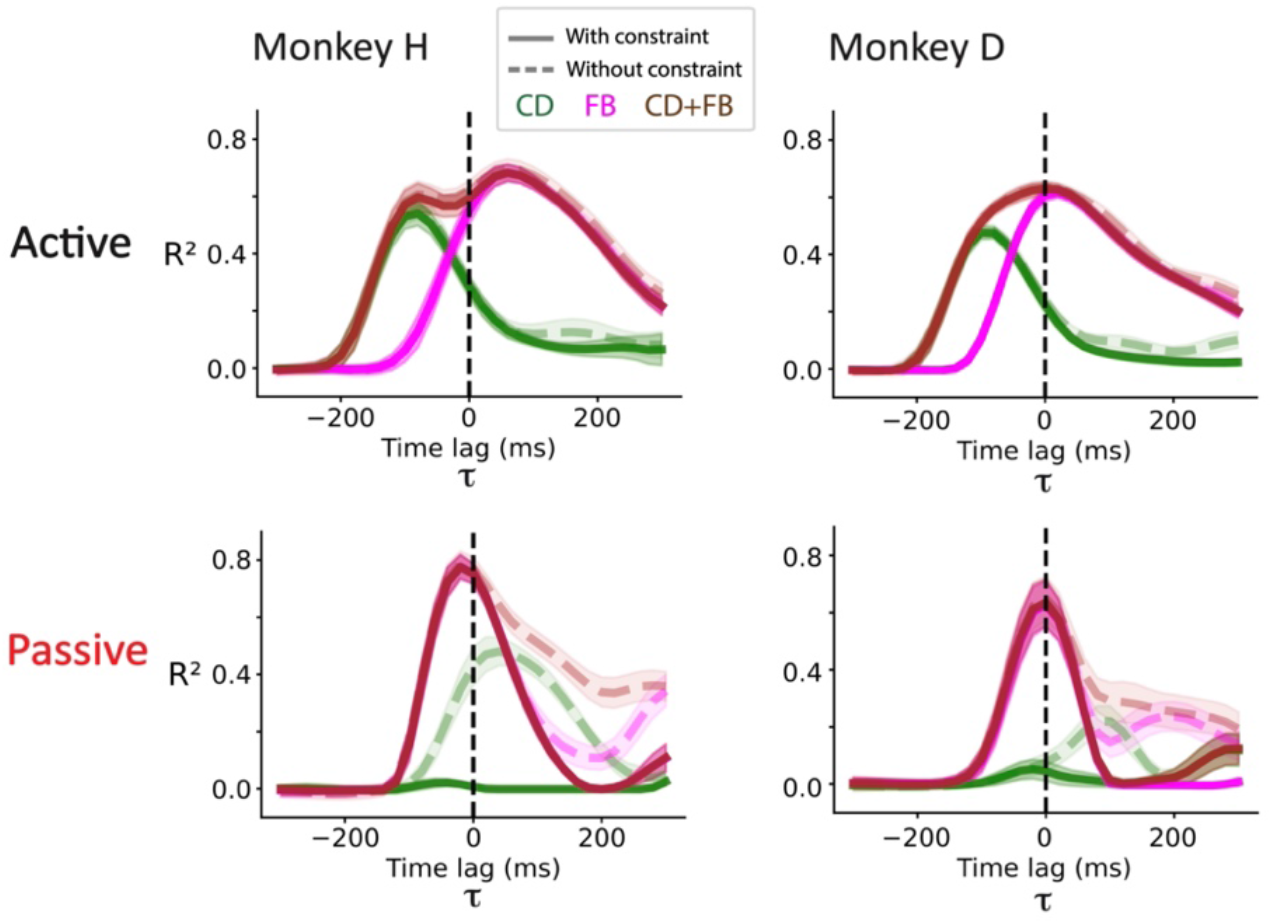
**Velocity decoding with or without constraining orientations of the signal dimensions**, expanding on results shown in **Figure 5B,C**. Removing the sign constraint (dotted lines) increased decoding accuracy mainly in passive trials, because the decoder could exploit return-related activity following the initial perturbation. Constraining the sign (solid lines) preserves the directional interpretation of the signal and prevents this return-related activity from inflating decoding performance, particularly for the corollary discharge signal.

**Figure 8.**
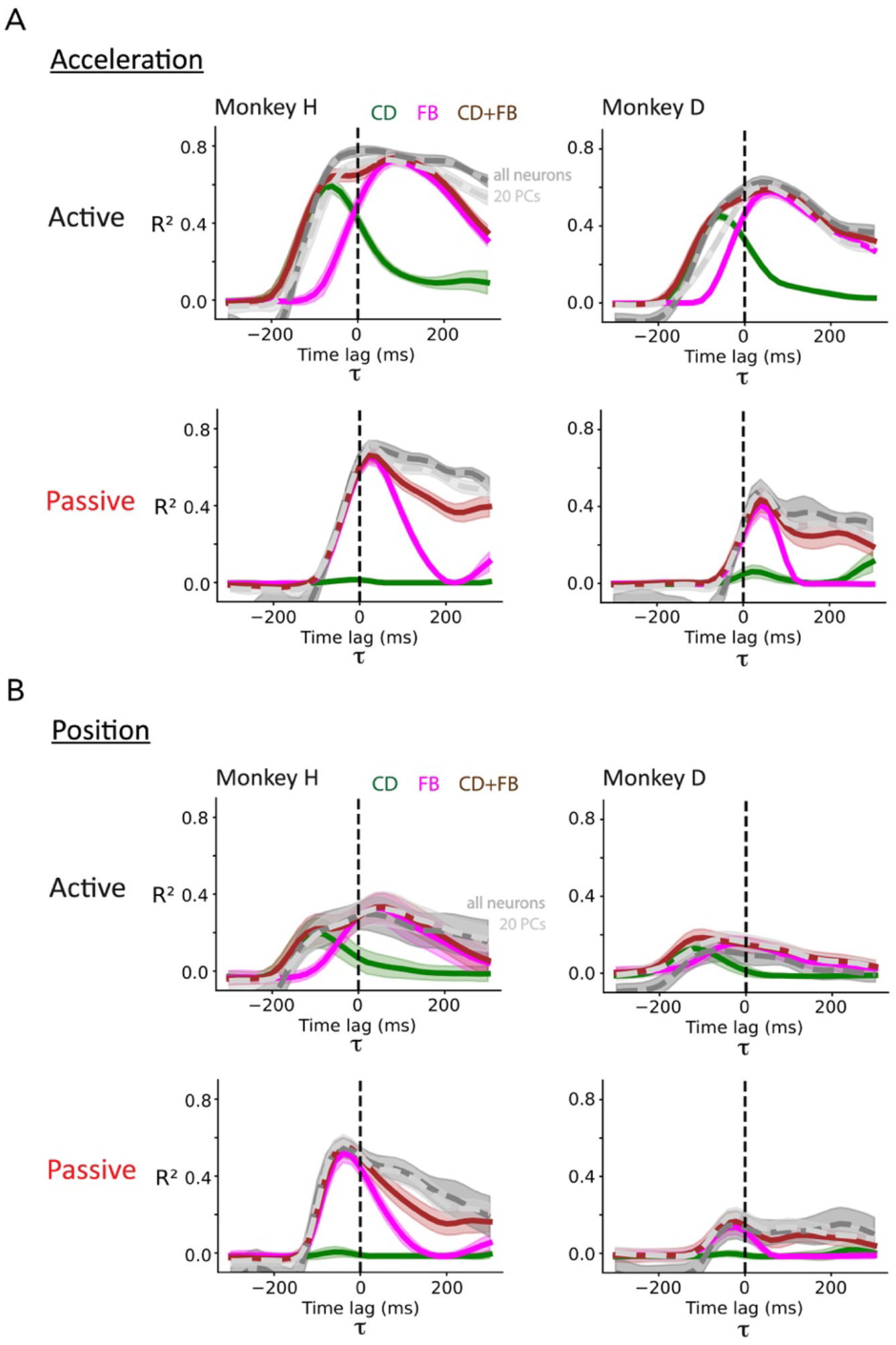
Full kinematic decoding in active and passive trials of Monkeys H and D. **A**. Acceleration decoding analysis. **B**. Position decoding analysis.

**Figure 9.**
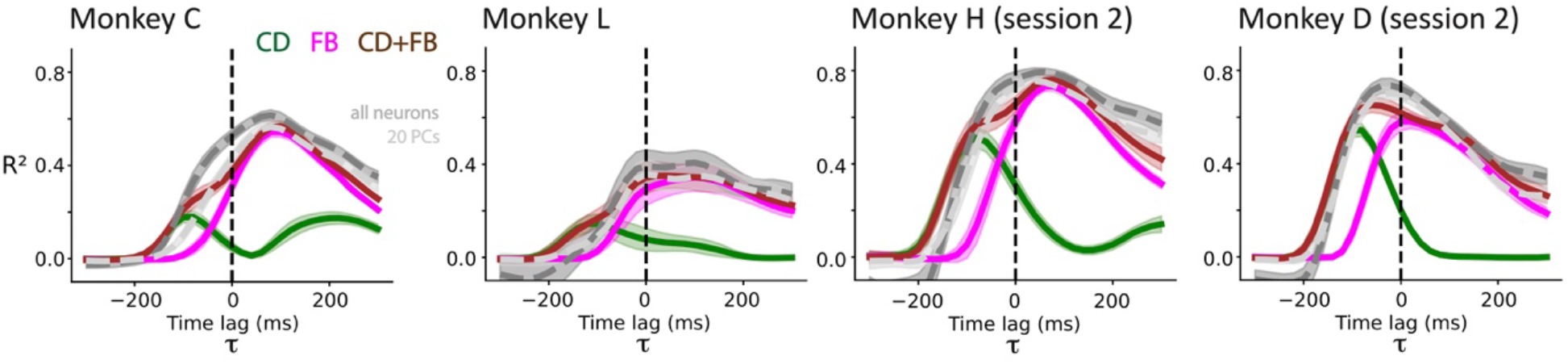
Effect of signal integration on estimating body state in Monkeys C and L and second sessions of Monkey H and D. Same format as Figure 5B. We achieved overall lower velocity decoding accuracy in Monkeys C and L, consistent with our observations that weak signals were collected from those two animals throughout analyses.

**Figure 10.**
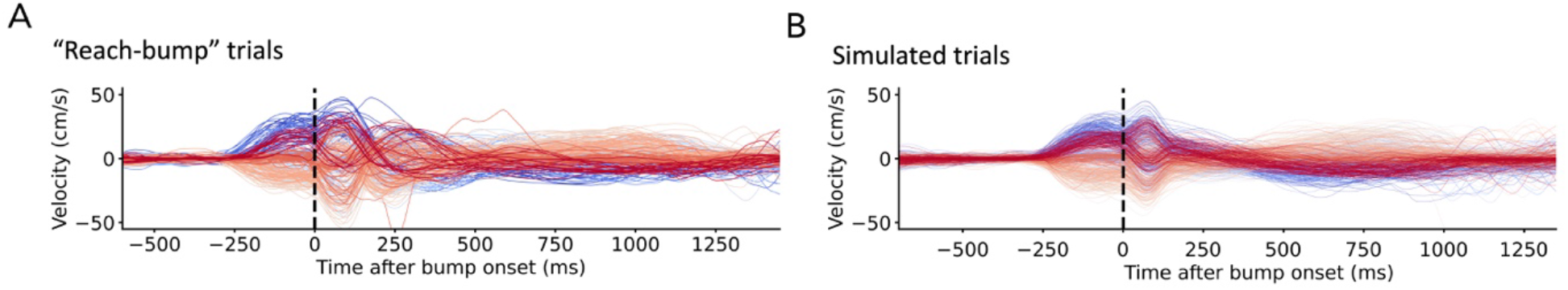
Experimental and simulated reach-bump kinematics. **A**. Hand velocity aligned to bump onset from experimentally-recorded trials. **B**. Hand velocity from simulated reach-bump trials by adding a Gaussian pulse to reach velocity in active trials. The resulting velocity profile closely matched those observed in the experimental trials.

**Figure 11.**
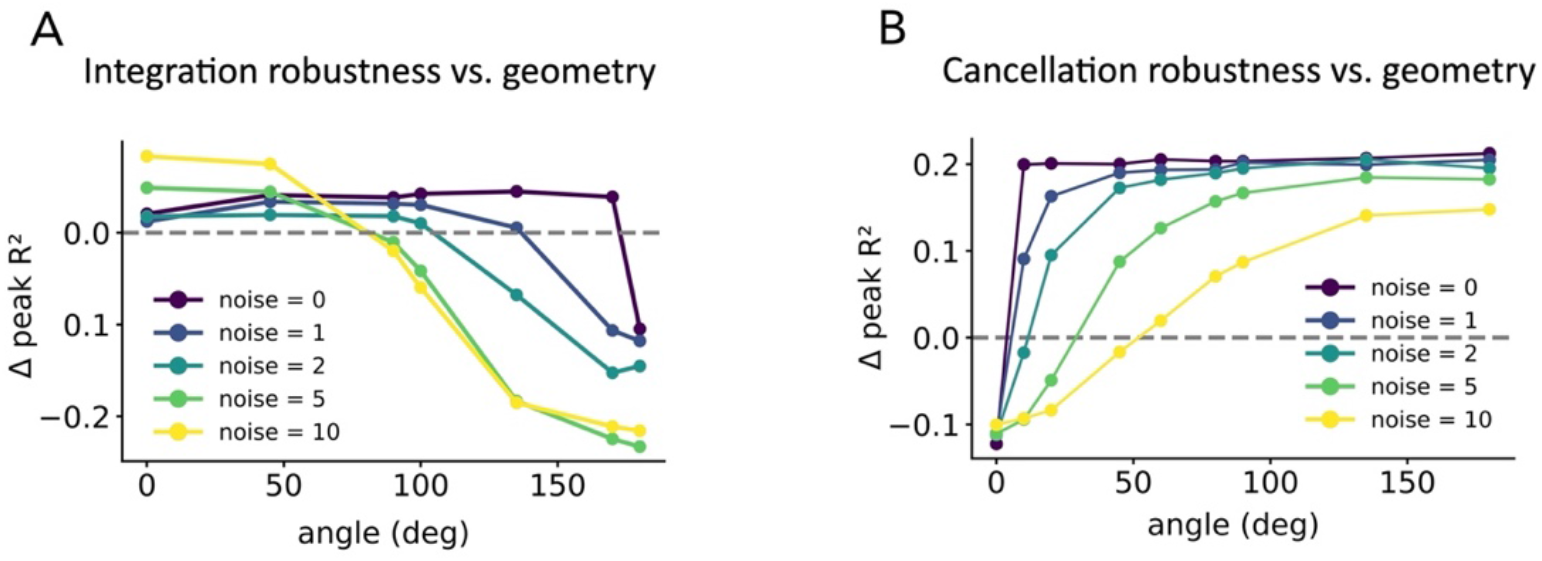
Simulation-based effects of representational geometry on decoding performance. **A**. Change in peak decoding accuracy (ΔR^2^) for combining corollary discharge and feedback signals relative to feedback alone as a function of the angle between them, for state estimation. At zero noise, combining the two signals improves decoding accuracy for any intermediate angle except for strictly opposite signals (180°). As system noise increases, geometries approaching opposition (∼180°) no longer improve decoding accuracy. **B**. Same analysis for perturbation classification. At zero noise, combining the two signals improves decoding accuracy for any intermediate angle. As system noise increases, geometries approaching alignment (∼0°) no longer improve decoding accuracy. Near-orthogonal geometry most robustly supports both functions across noise levels.

## Notes

### Competing Interest Statement

The authors have declared no competing interest.

### Summary of Updates

Abstract updated; Supplemental figure order updated

